# Elevated glucose in kidney organoids induces tissue-intrinsic inflammation driving epithelial detachment

**DOI:** 10.1101/2025.09.05.669416

**Authors:** Giulia Spennati, Mary C. Regier, Heather H. Ward, Vivek Das, Anil Karihaloo, Benjamin S. Freedman

**Affiliations:** Division of Nephrology; Department of Medicine; Institute for Stem Cells and Regenerative Medicine; Kidney Research Institute; University of Washington, Seattle, WA; Type 1 Diabetes and Kidney Disease, Global Drug Discovery, Novo Nordisk Research Center Seattle, Inc., Seattle, Washington; Immunobiology, Global Drug Discovery, Novo Nordisk, Lexington, Massachusetts; Kidney Biology, Global Drug Discovery, Novo Nordisk A/S, Maløv, Denmark; Novo Nordisk A/S, Soborg, Denmark

## Abstract

Diabetic kidney disease involves hyperglycemia, inflammation, and epithelial cell dysfunction, but the relationship between these factors is not well understood. We demonstrate that human kidney organoids treated with elevated glucose exhibit a phenotype of epithelial detachment driven by tissue-intrinsic upregulation of proinflammatory cytokines, which can be targeted therapeutically. High glucose induces morphological deterioration of organoids featuring podocyte and tubular cell detachment without cytotoxicity. Cytokine addition sensitizes organoids to intermediate concentrations of elevated glucose. Transcriptomic analysis reveals that high glucose levels affect cytokine, inflammation, signaling, and cell adhesion pathways, resembling changes in human diabetic kidneys. Inhibitors of cytokines and signaling pathways rescue the high glucose phenotype, which is independent of osmotic effects. Thus, elevated glucose triggers a tissue-intrinsic inflammatory cascade to produce an organ-specific phenotype in epithelial cells. This paradigm is relevant for understanding and potentially treating diabetic complications in the kidneys and possibly other organs.

## INTRODUCTION

Diabetes is currently the ninth cause of death worldwide affecting 537 million people. The hallmark of diabetes is elevated levels of blood glucose, with > 7 mM indicating severe disease. The acquisition of insulin resistance, as well as secondary complications of diabetes affecting various organs, are associated with major systemic changes in metabolism and inflammation, although the relationship between these factors is not fully understood^1^.

Amongst the organ-specific complications of diabetes, 40% of patients develop diabetic kidney disease (DKD), one of the leading causes of chronic kidney disease worldwide^2^. The pathogenesis of DKD is linked to inflammation, altered metabolism, and hemodynamic changes^3^. Proteinuria and albuminuria are common features, suggesting that impairments in the renal glomerular filtration barrier may be one of the primary initiators of the disease^3,4^. Podocytes, which are specialized epithelial cells that wrap around glomerular capillaries to establish the filtration barrier physically, may respond to stress by rearranging their actin cytoskeleton, which is hypothesized to contribute to DKD phenotypes ^5,6^. While substantial progress has recently been made in developing therapeutics to slow the progression of DKD, the significant residual risk for disease progression remains^7,8^.

Limited models are available for studying DKD, which has hampered our understanding of the disorder and ability to develop new therapies. Atlases of gene expression in DKD kidneys are emerging but have not yet been fully analyzed or interpreted ^7,9,10^. Rodent models in which hyperglycemia and obesity are induced can exhibit signs of glomerular injury, but these may not fully recapitulate the phenotypes or mechanisms of human DKD, due to species-specific differences^4^. Primary or immortalized renal cell cultures fail to replicate the complex structure of the nephron and are often dedifferentiated, which constrains their utility for studying this disorder^11^. Creating an in-vitro model that harbors multiple renal cell types organized in their natural complex arrangement becomes an attractive option to fill in this gap.

Induced pluripotent stem (iPS) cells can differentiate into a wide variety of human lineages and are a powerful tool for both disease modeling and regenerative medicine, relevant to diabetes and its complications^12,13^. Kidney organoids from iPS cells arise via a process of directed differentiation that mimics embryonic development^14–16^. Such organoids include well-differentiated epithelial cell types in contiguous, nephron-like segments, including podocytes, proximal tubules, and distal tubules, in addition to stromal and other cell types. Because organoids are grown in a petri dish, they are amenable to a wide range of experimental perturbations that would be difficult to achieve in a living organism.

We have previously described a transcriptomic signature of glomerular disease in our organoids under standard conditions, which may reflect the increased concentrations of glucose in our maintenance media relative to normal blood^17^. Oscillating treatments of 5 mM and 25 mM glucose in kidney organoids grown in suspension produced increased collagen deposition and susceptibility to SARS-CoV-2 infection, although changes to podocytes were not observed^18^. Similar to kidney tissue *in vivo,* organoid epithelial cells rapidly absorb glucose, a process that can increase cystogenesis in organoids with polycystic kidney disease mutations^19^.

In endothelial organoids, a combination of high glucose and inflammatory cytokines can produce a vasculopathy phenotype ^20^. Whether a DKD phenotype can be similarly recapitulated in kidney organoids is not yet known. Here, we conducted a systematic investigation of the effects of high glucose and cytokines on kidney organoids, with the goal of understanding what features of DKD these organoids might recapitulate.

## RESULTS

### High glucose alters the morphology of human organoids without overt toxicity

To model the effects of high glucose on renal cell types, human kidney organoids were differentiated in 11 mM glucose (the standard condition) and subsequently cultured for six days with concentrations of 5.5, 11, 33, and 100 mM glucose (**Figure 1A**). Organoids remained well preserved in glucose concentrations up to 11 mM by phase contrast microscopy, whereas organoids cultured in 33 and 100 mM glucose showed degradation of epithelia reflected as a ∼30% reduction in overall organoid number between days 3 and 6 (**Figures 1B-D and S1A**).

**Figure 1.**
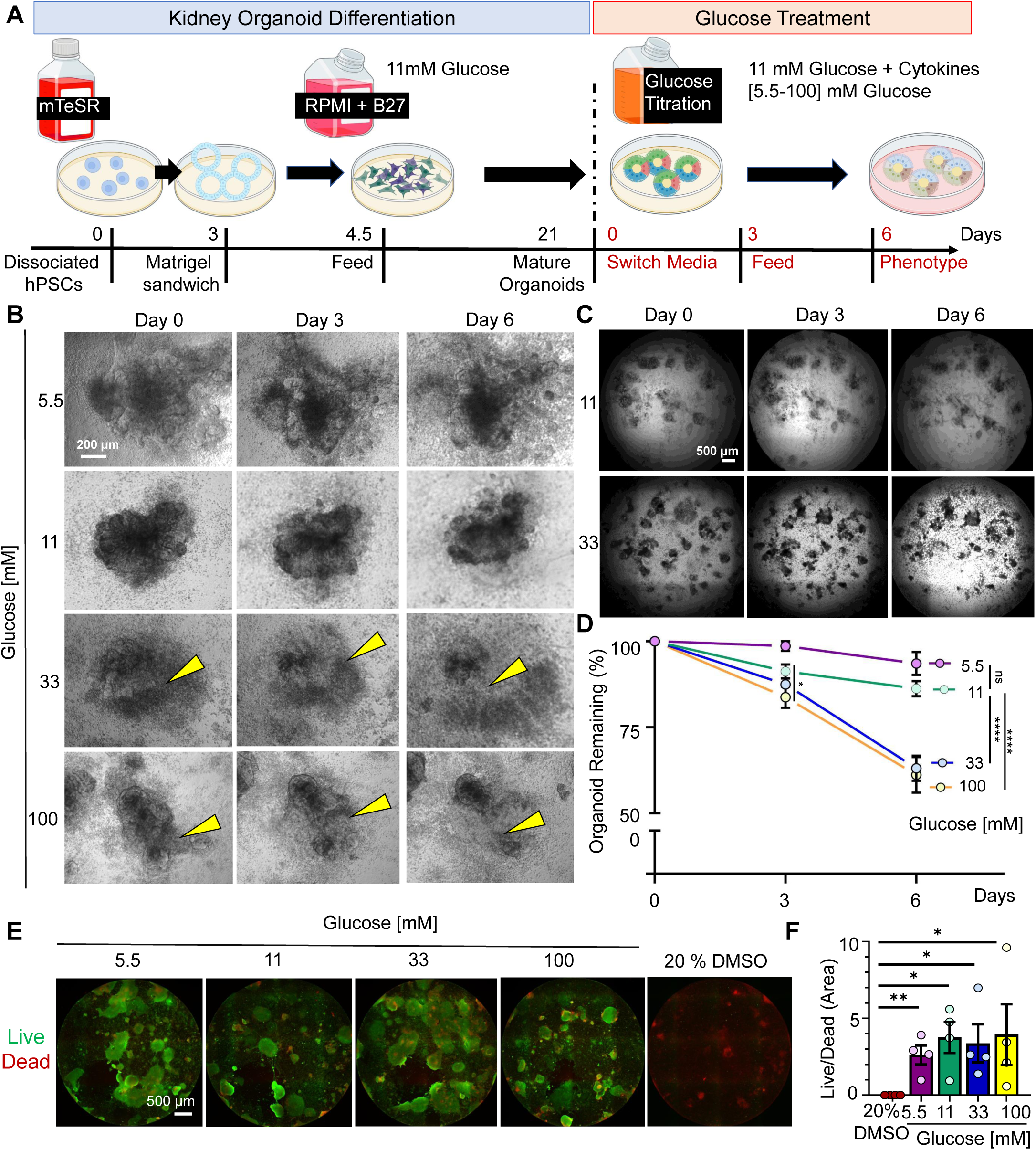
High glucose alters the morphology of human kidney organoids. A) Schematic of experimental workflow. Organoids are initially differentiated in 11mM glucose. On day 21, they are switched into media with different glucose concentrations. B) Time-lapse phase contrast images showing the progression of human kidney organoid morphology over six days of culture under four glucose concentrations. C) Time-lapse phase contrast images showing the progression of the whole plate over six days of culture, 11mM vs 33mM Glucose. D) Quantification of the percentage of organoids remaining on days 3 and 6 of glucose treatment, relative to day 0 (mean ± stderr from n = 8 independent experiments; **P<0.00, **P<0.01, *P<0.05 by one-way ANOVA). E) Wide-field images of the whole 96 wells showing live (green) and dead (red) assay. Positive control is 20% DMSO. f) Graph showing the ratio between live and dead areas (mean ± stderr from n = 4 wells pooled from 3 independent experiments; statistics as for (D).

To test whether high glucose was toxic to the organoids, we performed a live/dead assay at the day 6 endpoint. Results showed a prevalence of the green signal (live) over the red signal (dead) and similar ratios of live vs dead areas across glucose conditions (**Figure 1E-F**). A lactate dehydrogenase (LDH) assay performed on the media on days 3 and 6 also did not show significant toxicity in the higher glucose conditions (**Figure S1B**). Thus, the morphological changes observed in high glucose conditions were not due to overt cytotoxicity.

### High glucose causes podocyte detachment in kidney organoids

We investigated the impact of glucose on nephron segments in organoids. Interestingly, high levels of glucose (33, 100 mM) increased the number of small, single PODXL^+^ cells surrounding the main organoid body, suggesting detachment from the podocyte clusters that normally form in organoids (**Figure 2A**). To quantify, we segmented the total PODXL^+^ area into two sub-regions: intact area and detached area. The latter was further segmented into smaller regions of above-background intensity that were geometrically isolated from the podocyte clusters in the main organoid body (**Figure 2b**).

**Figure 2.**
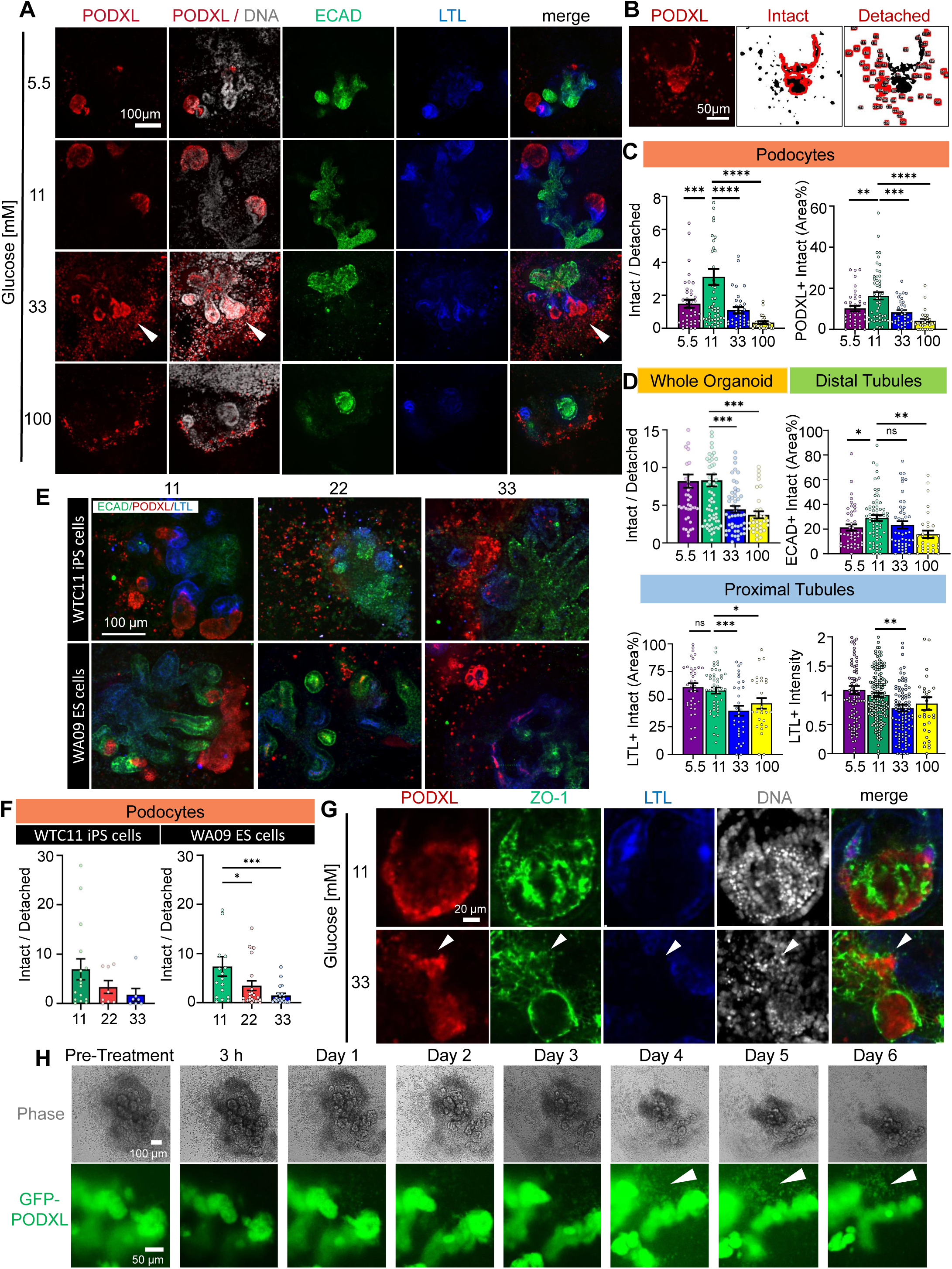
High glucose causes podocyte detachment in kidney organoids. A) Representative confocal immunofluorescence images showing a single optical section (20x magnification) for each condition. The white arrowhead indicates podocyte spreading. B) Representative image showing segmentation of intact and detached podocyte areas (outlined in red) based on raw immunofluorescence data (left). C) Quantification of detachment phenotype in podocytes (PODXL^+^) and D) whole organoids or tubular segments (mean ± stderr, n ≥ 30 organoids per condition pooled from 6 independent experiments; outlier data points are shown in fig. S2 ***P<0.00, **P<0.01, *P<0.05 by one way ANOVA). E) Representative confocal immunofluorescence images and F) quantification of WTC11 iPS and WA09 ES cells treated with 11, 22, or 33 mM glucose (mean ± stderr, n ≥ 8 organoids per condition pooled from 2 independent experiments; outliers data points are not displayed in the graphs, outlier data points are shown in fig. S2; ***P<0.00, **P<0.01, *P<0.05 by one-way ANOVA). G) Representative confocal immunofluorescence images showing a single optical section (20x magnification) for each condition. The white arrowhead indicates a flat sheet of podocytes adjacent to a larger aggregate. H) Representative time-lapse confocal immunofluorescence and phase contrast images of PODXL-GFP organoids.

Data indicates a decrease in the ratio of PODXL^+^ intact/detached area at 33 mM and 100 mM glucose concentrations and a 3-fold and 4-fold reduction of the PODXL^+^ intact area (**Figures 2C and S2A**). Results for the entire organoid epithelium (all three nephron segments combined, i.e., ECAD^+^, PODXL^+^, and LTL^+^ area) revealed a two-fold decrease in the intact/detached area ratio at the two higher glucose concentrations. Additionally, analysis of the nephron markers intact area reveals a 3-fold reduction in PODXL+ and LTL+ areas between 11 and 33 mM of glucose, and a non-significant reduction in the ECAD+ area (**Figures 2D and S2B-D**). Furthermore, glucose did not exhibit strong effects on PODXL intensity, suggesting that the cells remained differentiated, although a decrease in intensity was observed in LTL and ECAD, markers representing the other nephron segments (**Figures 2C and S2C-D**).

As 33 mM was relatively high for a physiological condition, we further investigated the intermediate concentration of 22 mM, and found that podocyte detachment was also observed with an intermediate level of severity, compared to 11 and 33 mM (**Figures 2E-F**). These findings were also reproduced in organoids derived from two different pluripotent stem cell lines, one male and one female, demonstrating that the effect was not unique to a particular line (**Figures 2E-F and S2E**).

PODXL^+^ cells in the detached areas expressed tight junctions, expressing *zonula occludens-1* (ZO-1) at the plasma membrane in a cobblestone pattern and contained intact, whole nuclei, indicating they were viable and remained epithelial cells (**Figure 2G**). To confirm the phenotype of podocyte detachment, organoids were differentiated from iPS cells encoding endogenous PODXL-tagged on its carboxyl-terminus with GFP^22^, and individual organoids were tracked over 6 days in 11 mM and 33 mM of glucose. We detected progressive outgrowth of PODXL^+^ cells in the regions surrounding podocyte aggregates in the main organoid body, with a substantial increase of these detached cells by day 4, contemporaneous with significant modifications of organoid morphology and area shrinkage (**Figures 2H, S2F, and Movies 1-2**). Thus, a major effect of high glucose was the induction of podocyte damage and detachment from larger aggregates.

### Inflammatory cytokines induce podocyte detachment in physiological glucose

To test whether these phenotypes could be observed under more clinically relevant glucose concentrations, we supplemented our media with TNF-alpha (tumor necrosis factor-alpha), a major cytokine observed in the diabetic milieu^20,23^, which is known to be upregulated in DKD in humans^24^. When 1 ng/ml of TNF-alpha was added to the growth media containing 5.5 mM, 11 mM, or 33 mM of glucose for six days, a reduction in organoid number was observed, which was more pronounced in 11 mM versus 5.5 mM glucose (**Figures 3A-B**). TNF-alpha did not cause increased cell death across glucose conditions as indicated by live/dead assay or LDH assay (**Figures 3C-D and S3G**).

**Figure 3.**
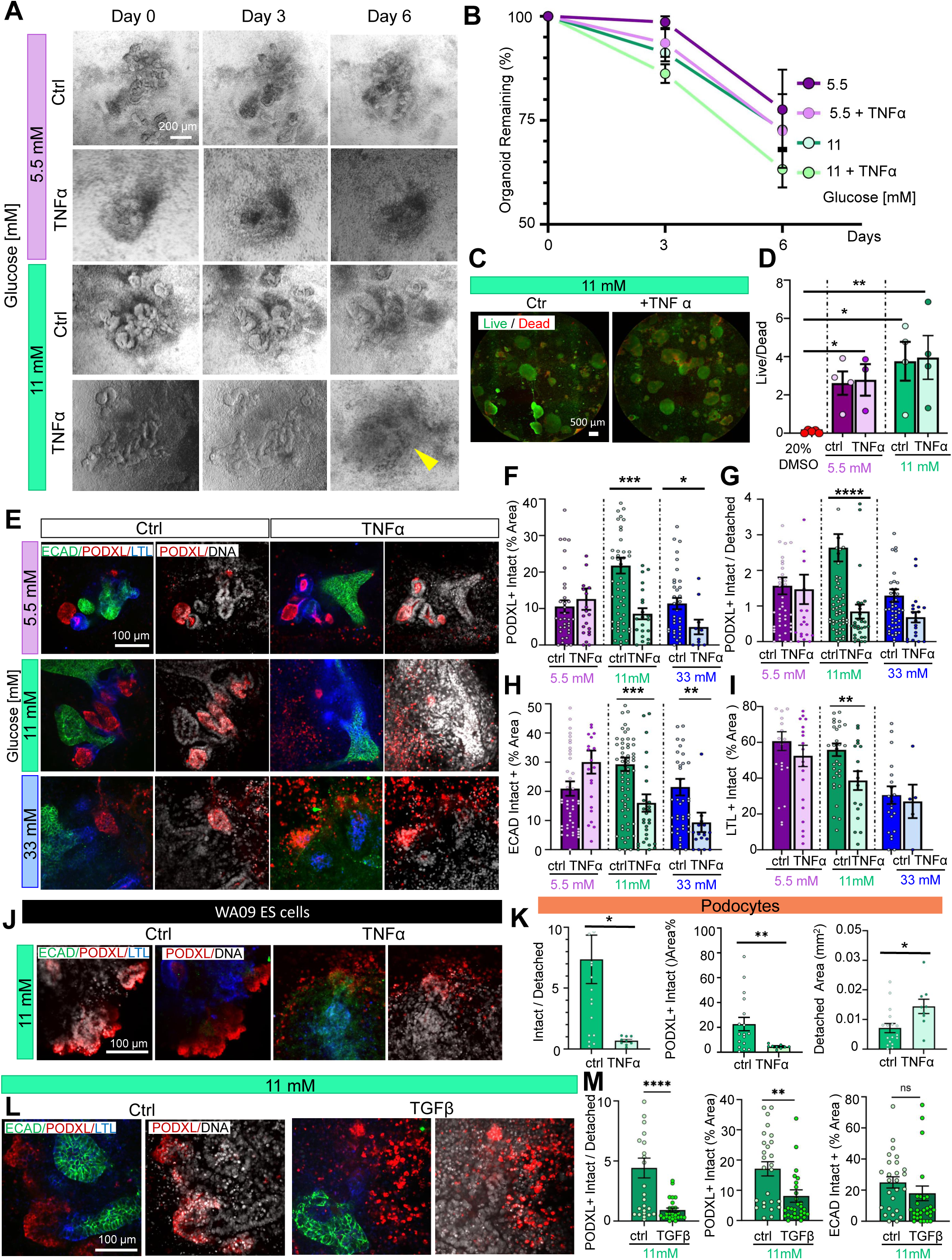
TNF-alpha and TGF-beta sensitize organoids to physiologically elevated glucose. A) Time-lapse phase contrast images showing the progression of human kidney organoid morphology over six days of culture. B) Quantification of the percentage of organoids remaining on days 3 and 6 of glucose treatment ± TNF-alpha, relative to day 0 (mean ± stderr from n ≥ 4 independent experiments). C) Wide-field images and D) quantification of live-dead staining in organoids (mean ± stderr from n = 4 wells pooled from 3 independent experiments). E) Representative confocal immunofluorescence images showing a single optical section (20x magnification) for each condition. F) Quantification of PODXL intact vs detached areas and G) PODXL, H) ECAD, and I) LTL intact areas (mean ± stderr, n ≥ 21 organoids per condition pooled n ≥ 6 independent experiments; ***P<0.00, **P<0.01, *P<0.05 by one way ANOVA. Outlier data points are shown in fig. S3). J) Representative confocal immunofluorescence images showing a single optical section (20x magnification) of organoids derived from WA09 ES cells ± TNF-alpha. K) Quantification of PODXL intact vs detached areas, % intact, and % detached areas (mean ± stderr, n ≥ 18 organoids per condition pooled n ≥ 2 independent experiments; outliers data points are shown in figure s-3, ***P<0.001, **P<0.01, *P<0.05 by one way ANOVA). L) Representative confocal immunofluorescence images showing a single optical section (20x magnification) for each condition. M) Quantification of PODXL intact vs detached areas and PODXL and ECAD intact areas (mean ± stderr, n ≥ 14 organoids per condition pooled from 3 independent experiments; outliers data points are shown in supplementary figure s-3, ***P<0.001, **P<0.01, *P<0.05 by one way ANOVA).

After TNF-alpha exposure, we observed an increase in the number of small, single PODXL^+^ cells surrounding the main organoid body, suggesting detachment from the podocyte clusters that normally form in organoids (**Figure 3E**). Strikingly, the PODXL^+^ intact vs. detached area ratio was significantly reduced when TNF-alpha was added to the 11 mM glucose condition, but not 5 mM or 33 mM glucose (**Figures 3F and S3A-B**). Intact areas of PODXL^+^, LTL^+^, and ECAD^+^ were all reduced when TNF-alpha was added to 11 mM glucose, with a minor impact on the other conditions (**Figures 3G-I and S3C-E**). Co-staining of tight junctions (ZO-1) with PODXL confirmed an increased number of podocytes outside of the glomerular ‘tuft’ when treated with TNF-alpha (**Figure S3F**).

Collectively, these experiments suggested that organoids undergo podocyte detachment when TNF-alpha is combined with levels of glucose that would be considered diabetic in a living person (11 mM). A comparable phenotype was observed in organoids derived from a second pluripotent stem cell line (**Figures 3J-K and Figure S3H**).

In addition to TNF-alpha, we also tested the effects of TGF-beta, which is another major cytokine associated with DKD^3,20^. Similar to TNF-alpha, TGF-beta exposure caused a four-fold reduction of PODXL^+^ intact vs. detached area ratio and a two-fold reduction of PODXL^+^ intact area, whereas ECAD^+^ area was unaffected (**Figures 3L-M and S3I**). These findings suggested that both TNF-alpha and TGF-beta can contribute to podocyte damage at physiologically relevant glucose concentrations.

### scRNA-seq reveals high glucose induces cytokines and signaling

To further identify mechanisms that might mediate the effects of high glucose, we performed an unbiased single-cell RNA sequencing analysis of kidney organoids cultured in 11 mM vs. 33 mM glucose for 6 days. Cells were sequenced and grouped into eleven major clusters, including epithelial, stromal, and proliferative (**Figures S4A-C**). We focused on the epithelial cluster and identified eight sub-clusters, including podocyte, proximal tubular, and distal tubular segments, as well as early glomerular and stroma-like epithelial cells, with all of these cell types represented in both of the glucose treatment conditions (**Figures 4A and S4D-E**).

**Figure 4.**
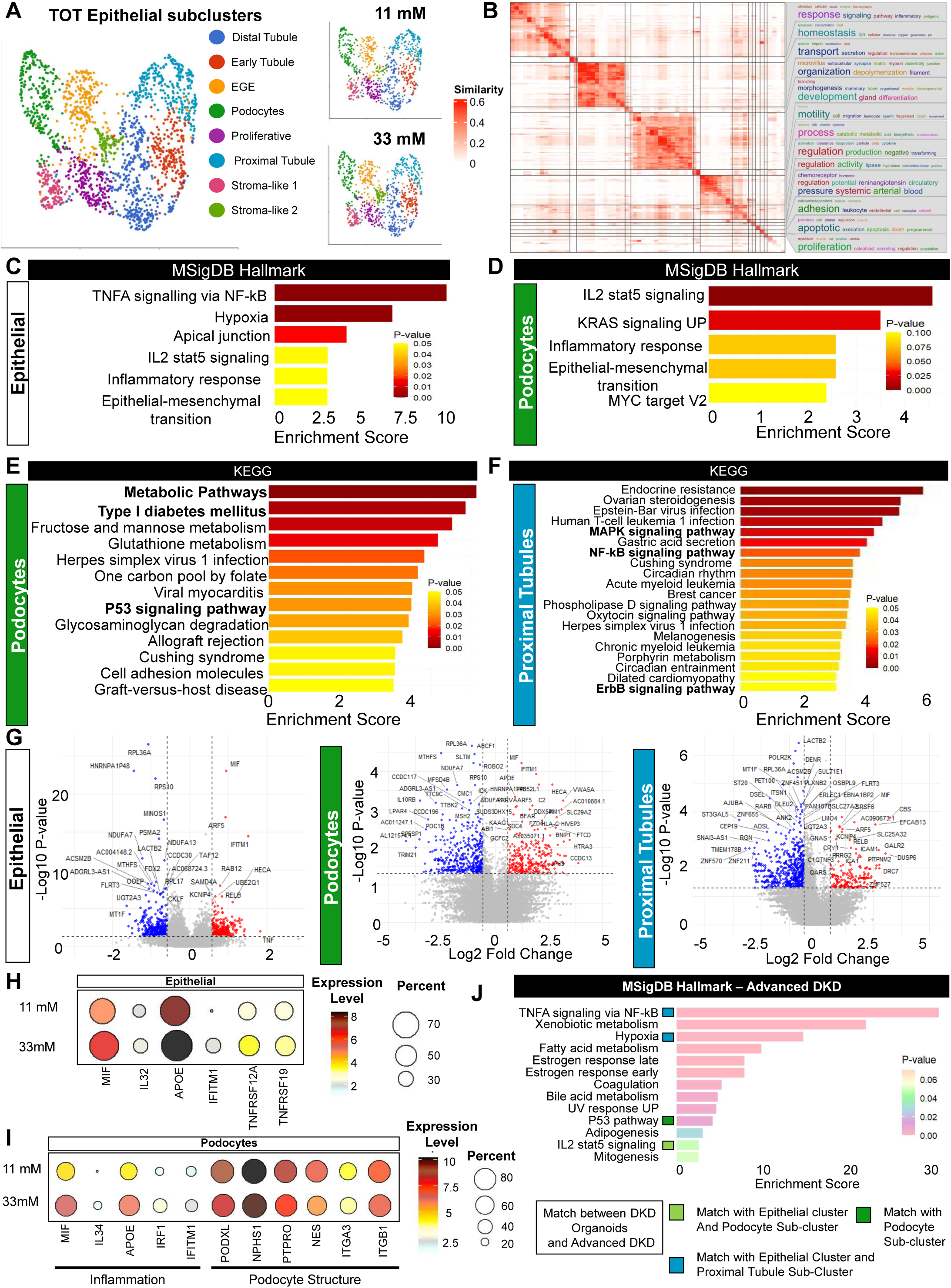
scRNA-seq analysis reveals that high glucose induces cytokine and signalling pathways. A) UMAPs showing the epithelial cluster, with sub-clusters labeled by cell type. B) GO term semantic similarity for the epithelial cluster. C) Top upregulated pathways with MSigDB Hallmark dataset in the epithelial cluster, and D) the podocyte subcluster. E) Top upregulated pathways with the KEGG dataset in the podocyte subcluster and F) the proximal tubule subcluster. G) Volcano plots showing up- and down-regulated genes in 33 vs. 11 mM glucose, with cutoffs of fold change > 1.5 and P value < 0.05 in the epithelial cluster, podocyte subcluster, and proximal tubule sub-cluster. H) Expression of genes of interest between 11 vs 33 mM of glucose in the epithelial cluster and I) the podocyte sub-cluster. J) Enriched pathways in the human advanced DKD cohort identified with gene set enrichment analysis using the MSigDB Hallmark dataset, indicating pathways in common with both the epithelial cluster and the podocyte sub-cluster in the organoid dataset (light green), both the epithelial cluster and proximal tubule sub-cluster (blue), and only with the podocyte sub-cluster (dark green).

We performed gene set enrichment analysis to identify enriched pathways in the epithelial cluster as a whole, as well as its podocyte and proximal tubule sub-clusters. Semantic similarity analysis of Gene Ontology (GO) terms in 33 mM vs. 11 mM glucose produced functional summary words, including organization, regulation, motility, and adhesion in the epithelial cluster; organization, differentiation, regulation, and response in the podocyte sub-cluster; and metabolic process, proliferation and transport in the proximal tubule sub-cluster (**Figures 4B and S5A-B**).

MSigDB (Molecular Signatures database) Hallmark analysis identified TNF-alpha/NF-kB (nuclear factor kappa-light-chain-enhancer of activated B cells) signaling as a top-upregulated pathway in the epithelial cluster and proximal tubule sub-cluster, along with inflammation-related pathways and epithelial-to-mesenchymal transition, which were also highly enriched in the podocyte sub-cluster (**Figures 4C-D and S5C-D**)^25^. Additionally, KEGG (Kyoto Encyclopedia of Genes and Genomes) analyses identified upregulation of metabolic pathways, P53 signaling ^4,26^, and type 1 diabetes in the podocyte sub-cluster; type II diabetes in the epithelial cluster; and NF-kB, MAPK, and ErbB signaling pathways in the proximal tubules (**Figures 4E-F and S5D-E**).

Comparing 33 mM glucose to 11 mM, MIF (macrophage migration inhibition factor) emerged as the top-upregulated gene in the complete dataset, the epithelial cluster, and the podocyte sub-cluster (**Figures 4F-G and S5D**). MIF binds receptor CD74 (cluster of differentiation 74), leading to downstream signaling through MAPK, PI3K/AKT (phosphoinositide-3-kinase–protein kinase B/Akt), or NF-kB pathways.^27,28^ We identified 310 upregulated and 399 downregulated genes in the epithelial cluster, including TNF and other genes linked with inflammation, and observed upregulation of MIF in the epithelial cluster and both the podocyte and proximal tubule sub-clusters (**Figures 4 G-H**). Within the podocyte sub-cluster, we observed a similar upregulation of inflammatory genes, along with the downregulation of podocyte structural markers (**Figure 4I**)^29,30^.

To further validate our analysis, we compared our scRNA-seq data with a human kidney biopsy bulk sequencing dataset from 22 advanced DKD patients and 9 controls^10^. A gene set enrichment analysis using the MSigDB Hallmark database between the organoid model and the DKD population identified several common pathways, including TNF-alpha/ NF-kB, hypoxia, P53, and IL2 stat5 signaling pathway (**Figure 4J**). Of 56 differentially expressed genes (22% of all analyzed) appearing in both the 33 mM glucose organoid condition and DKD biopsy samples, 36 genes (64 %) showed similar directionality of differential expression in both datasets, with 27% upregulated and 37% downregulated (**Figures S5F-I**).

We surveyed the broader literature to determine whether these pathways were previously identified in analyses of DKD tissue. The European Renal cDNA bank (ERCB) consortium linked DKD progression to NF-kB-driven inflammation pathways^31^. In our scRNA-seq dataset, upregulation of the MSigDB Hallmark gene set term “TNF-alpha signaling via NF-kB,” as well as the individual genes *TNF, TNFRSF12A*, and *TNFRSF19*, was particularly interesting, because MIF is a proinflammatory cytokine believed to activate TNF-alpha^32,33^, a key player in CKD development^23^. Prior studies reported the enrichment of cell adhesion and inflammatory response pathways in DKD tissue, including cytokine-cytokine receptor interaction (glomeruli) and MAPK (tubulointerstitium)^34^. Thus, our scRNA-seq analysis suggested that high glucose levels promote TNF-alpha and MIF expression, driving inflammation and podocyte injury.

### Inhibitors protect organoids from the effects of high glucose

We further investigated the increased expression of MIF and TNF-alpha as a potential pathway in this process. Immunoblot analysis confirmed increased MIF expression at 33 mM of glucose (**Figures 5A and S6A**). In our scRNA-seq dataset, MIF expression was highest in the podocytes, amongst the sub-clusters of the epithelial cluster (**Figure 5B**). *TNF* expression was increased ∼5-fold in high glucose by qPCR analysis (**Figure 5C**). Thus, upregulation of these related cytokines appeared reproducible in our follow-up analyses.

**Figure 5.**
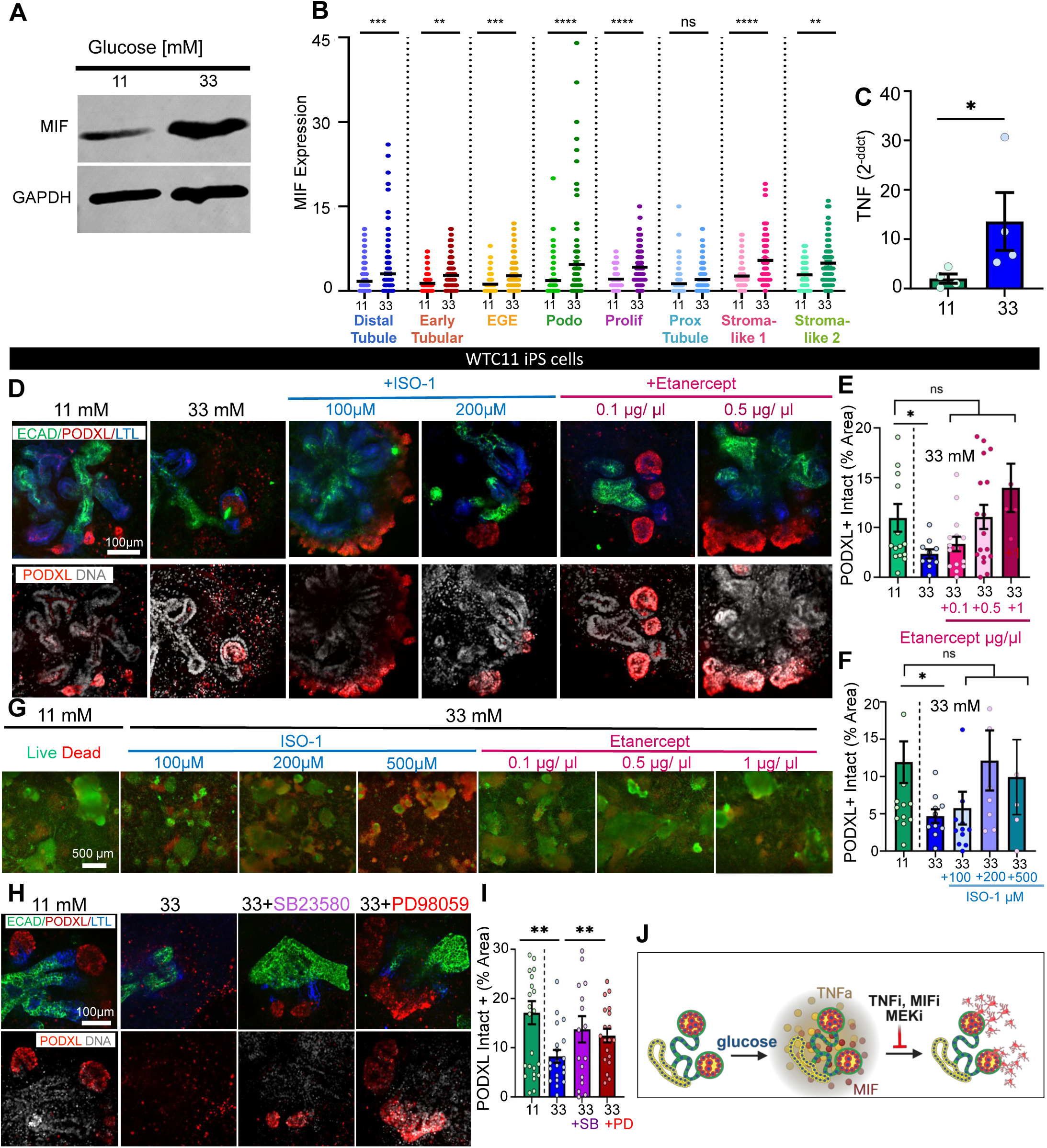
MIF or TNF-alpha inhibition protects organoids from the effects of high glucose. A) Representative MIF immunoblot with GAPDH loading control. B) Expression of MIF within different epithelial sub-clusters in 11mM vs 33mM analyzed with scRNAseq. C) qPCR analysis of TNF expression comparing 33mM with 11 mM of glucose (mean ± stderr, n =4 independent experiments; *P<0.05 by student’s t-test). Outlier data points are shown in Figure S4. D) Representative confocal immunofluorescence images of kidney organoids in different treatment conditions showing a single optical section (20x magnification). E) Quantification of PODXL intact area ± ISO-1 and F) ± etanercept (mean ± stderr, n ≥ 5 organoids per condition pooled n = 2 independent experiments; outlier data points are shown in figure S6. ***P<0.00, **P<0.01, *P<0.05 by one way ANOVA). G) Representative wide-field images of whole 96 wells showing live (green) and dead (red) assay for ISO-1 and etanercept. H) Representative confocal immunofluorescence images showing a single optical section (20x magnification) with I) quantification of PODXL intact area and intact vs detached areas ± SB23580 or ± PD98059 (mean ± stderr, n ≥ 19 organoids per condition pooled from n =3 independent experiments). Outlier data points are shown in fig. S6. ***P<0.00, **P<0.01, *P<0.05 by one-way ANOVA). J) Schematic diagram representing the model of injury. Kidney organoids are cultured in high glucose, which leads to tissue-intrinsic expression of inflammatory cytokines (MIF and TNF-alpha), and consequent detachment of podocytes and other epithelial cells. Inhibition of TNF-alpha, MIF, or MEK protects the epithelial cells from detachment.

To further validate these findings and investigate possible therapeutic strategies, kidney organoids were cultured with 33 mM glucose with inhibitors of either TNF-alpha (etanercept) or MIF (ISO-1) for six days. Both etanercept and ISO-1 prevented the podocyte injury phenotype in the 33 mM glucose treatment condition (**Figures 5D-F and S6C-F**). No toxicity was observed by live/dead assay up to 200 μM of ISO-1 and 0.5 µg/μl of etanercept (**Figure 5G**). Comparable results are identified in a second cell line (**Figures S6E-F**).

As our scRNA-seq showed upregulation of MAPK in proximal tubules, and this pathway is generally linked with inflammation^35^ we tested the effect of two MAPK inhibitors, SB203580 (an ATP-competitive inhibitor of p38 MAPK) and PD98059 (an ERK1/2 inhibitor). Each of these inhibitors exhibited a protective effect on podocyte morphology when added to the media with 33 mM glucose (**Figure 5H**). The intact PODXL area for each of these was partially rescued towards the values obtained with 11 mM glucose (**Figures 5H-I and S6G-H**).

Collectively, these studies suggested a model in which high glucose induces inflammation and increased cytokine expression (MIF and TNF-alpha). Subsequent detachment of podocytes and other epithelial cells from the main organoid body (**Figure 5J**).

### Deterioration in high glucose is not due to osmotic lysis

To test whether osmotic effects of high glucose were involved in the observed phenotype, organoids were cultured with increasing concentrations of mannose versus equimolar glucose or a combination of the two sugars. Mannose is an isomer of glucose with equal molecular weight and osmotic characteristics that can be absorbed by human cells but is not metabolized in the same way, providing a control for osmotic effects.^36,37^ Kidney organoids cultured with 33 mM mannose or combined sugars (11 mM glucose + 22 mM mannose) did not show signs of podocyte detachment, in contrast to 33 mM glucose alone (**Figures 6A-B and S6I-J**).

**Figure 6.**
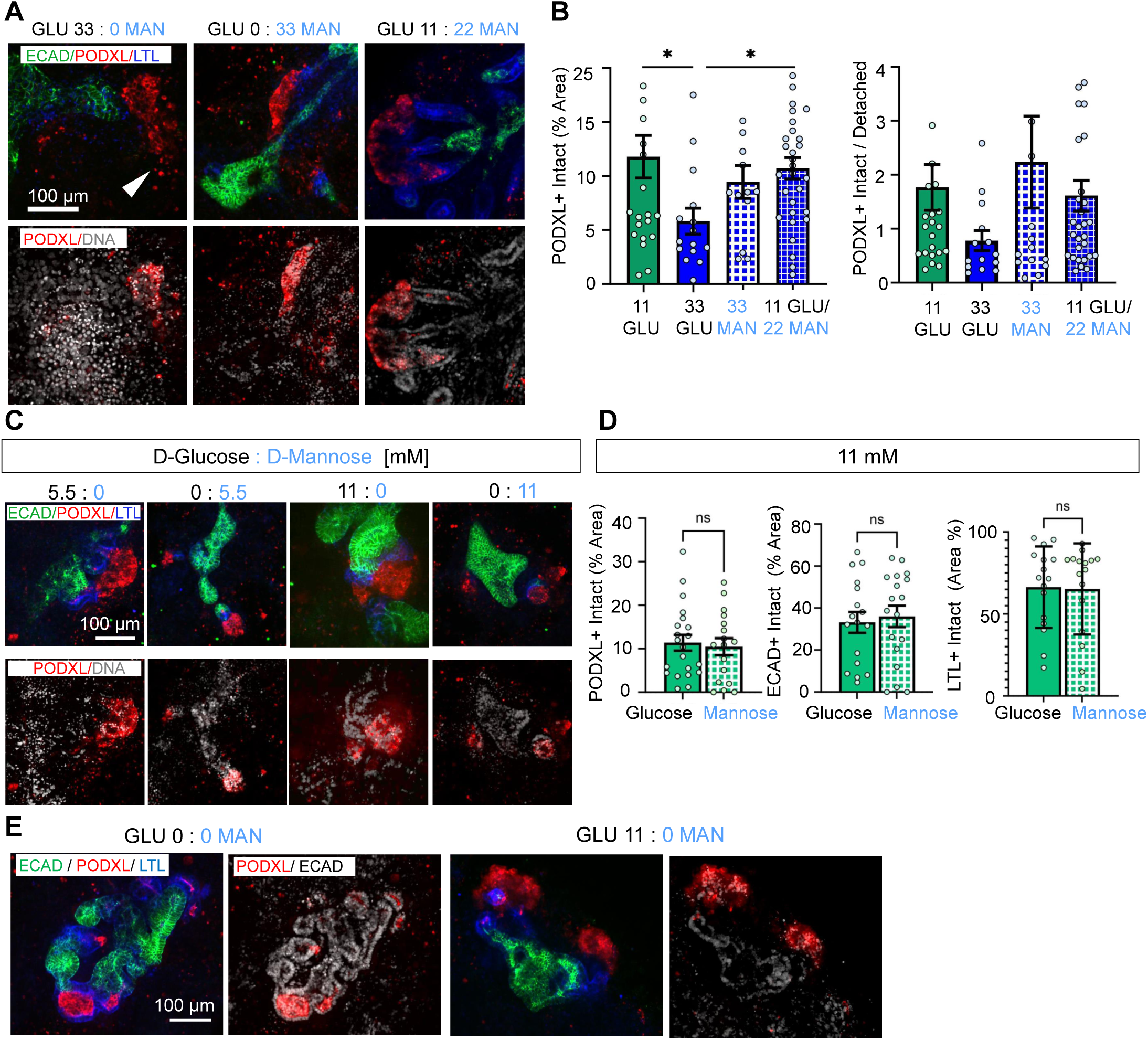
Deterioration in high glucose is not due to osmotic lysis. A) Representative confocal immunofluorescence images showing a single optical section (20x magnification) comparing high glucose, mannose, or their combination. The white arrowhead indicates podocyte spreading. B) Quantification of PODXL^+^ areas and intact vs. detached ratio (mean ± stderr, n ≥ 12 organoids per condition pooled from 4 independent experiments; outlier data points are shown in figure S6; ***P<0.00, **P<0.01, *P<0.05 by one-way ANOVA). C) Representative confocal immunofluorescence images showing a single optical section (20x magnification) comparing equimolar concentrations of glucose and mannose. D) Quantification of PODXL, ECAD, LTL areas comparing 11 mM glucose vs mannose (mean ± stderr, n ≥ 15 organoids per condition pooled from 5 independent experiments; outlier data points are shown in fig S6 ***P<0.00, **P<0.01, *P<0.05 by one way ANOVA). E) Representative confocal immunofluorescence images showing a single optical section (20x magnification) comparing 0 vs 11 mM glucose with no mannose.

At lower concentrations of glucose or mannose (5.5 mM and 11 mM), organoids showed comparable morphology across the three nephron segment markers (**Figures 6C-D**). Surprisingly, the morphology of the kidney organoids appeared well-maintained after six days of culture, even in the complete absence of glucose or mannose (**Figure 6E**). These data suggested that the observed effects of high glucose were not due to osmotic stress.

## DISCUSSION

### A new experimental model for glucose effects

Kidney organoids have emerged as valuable models of renal disease, but their response to high glucose and cytokines has not been studied in detail. Our experiments identify an injury phenotype under high glucose concentrations (>22 mM), in which podocytes detach from the main organoid body. This phenotype significantly resembles findings from analyses of biopsies^6,38^ and urine samples from DKD patients,^39,40^ and of animal models of DKD,^38,41^ in which podocyte loss and detachment from the glomerular basement membrane, rather than apoptosis, was an early step associated with pathophysiology. Many genes or pathways differentially expressed in organoids in high glucose are similarly altered in human kidneys from patients with DKD, supporting the relevance of this new model^10^.

In contrast to this study, a previous report did not describe the spreading of podocyte clusters in kidney organoids grown in suspension and treated with oscillating levels of 25 mM and 5 mM glucose^18^. It is possible that the injury phenotype was not detected in that study because it was focused on SARS-CoV-2 infection in tubular cells^18^, or because the smaller, adherent organoids in our differentiation protocol may be advantageous for eliciting or detecting podocyte spreading^14^. This mirrors the situation for experimental models of polycystic kidney disease, for which expression of the pathognomonic phenotype depends on organoid size and culture conditions^14,42^.

### Glucose triggers inflammation

This new model *in vitro* offers interesting insights into the pathophysiology of diabetes and its complications. Immune pathway activation has been associated with the development of insulin resistance and is suggested to play a role in diabetic complications in organs, including the retina, nervous system, and kidneys.^1,43^ How this works mechanistically is not well understood and can be difficult to deconvolve in model organisms wherein many different organ systems, including the immune system and adipose tissues, can be sources of cytokines.^1,9^ In our comparatively simpler human kidney organoids, which lack a functional immune system, high glucose induces an inflammatory response that contributes to the emergence of a specific cellular response appropriate to diabetes. In contrast to vascular organoids, where a combination of high glucose (75 mM) and cytokines is required to elicit a diabetic phenotype,^20^ in kidney organoids the addition of glucose alone (33 mM) is sufficient, thus the inflammatory response could be pinpointed as downstream of the glucose stimulation. Notably, mannose does not induce the same response as glucose, suggesting a possible role for metabolism in this process as opposed to osmotic effects.

Both scRNA-seq analysis and functional studies indicate that MIF and TNF-alpha play an important role in the phenotype of epithelial detachment observed in high glucose. These inflammatory cytokines have previously been associated with DKD in human and mouse models.^33,41,43^ High urinary levels of MIF have been observed in individuals with acute kidney injury (AKI)^44^ and type 1 diabetes,^45^ and several clinical studies have reported elevated levels of urinary cytokines, including TNF-alpha,^24^ in patients with proximal tubular damage damage^46^ or with diabetes.^47^ Kidney proximal tubules, via the sodium-glucose cotransporter 2 (SGLT2), are responsible for glucose reabsorption. The hyperglycemic state, due to diabetes, is thought to an energetic overload on the proximal tubules, leading to cellular stress and damage, which triggers cytokine release and inflammation.^35,48^ This paradigm for direct effects of glucose triggering the innate immune response might also be generalizable, for instance, to the acquisition of insulin resistance, extrarenal diabetic complications, or other disease states. Kidney organoids can respond to inflammatory cues to produce changes in gene expression and cell death relevant to COVID-19 and APOL1-mediated kidney disease.^22,49^

Among the different cell types in kidney organoids, podocytes substantially upregulate MIF in response to glucose, raising the possibility that cytokines are expressed by the same cells that respond to them. Epithelial cell detachment was observed in response to glucose not only in podocytes, but also in tubular cells. Our recent studies have also identified a role for glucose in augmenting the cystic phenotype in organoids with polycystic kidney disease mutations, which may suggest a common role for glucose in regulating epithelial structure, differentiation status, and activity.^19^ Notably, in the context of acute kidney injury *in vivo*, as arises due to rapid chemical or ischemic insult, there is also evidence that kidney epithelial cells respond by utilizing immune response pathways.^50–52^ The tendency of kidney epithelial cells to respond in this manner may underlie the sensitivity of the kidneys to diabetic complications compared to certain other organs.

### Clinical implications

The use of cytokine inhibitors to treat diabetes and diabetic complications remains an emerging area of investigation. Etanercept, the TNF-alpha-specific inhibitor used in our study, was initially assessed for safety properties in patients with kidney disease and does not show broad nephrotoxicity, although safety concerns remain regarding the potential side effects of this drug.^53,54^ ISO-1, a MIF inhibitor, has not yet been tried in humans, but in diabetic mice it significantly reduced blood glucose levels, albuminuria, extracellular matrix accumulation, and EMT.^41^ Pentoxifylline, a phosphodiesterase inhibitor that has both hemorheological and anti-inflammatory properties, has shown promise in smaller clinical trials for DKD and is currently being studied in a larger cohort.^55,56^ MIF inhibitors are in development for the clinic, including an ongoing stage II trial of ibudilast, a phosphodiesterase inhibitor with effects on MIF, for amyotrophic lateral sclerosis,^57^ but have not yet been applied in patient populations with diabetes. Our findings support continued clinical investigation of cytokine inhibitors for DKD. With regard to downstream signals such as MAPK, agonists of the beta 2 adrenergic receptor, which can regulate this pathway, have recently been shown to be protective in CKD patients, with supporting data showing a reversal of phenotype in a DKD mouse model^58^. Current therapies such as SGLT2 inhibitors and glucagon-like peptide-1 agonists may also affect cytokine responses.^8,59^

A limitation of our study is that, in the absence of TNF-alpha, relatively high levels of glucose (≥ 22 mM) are needed to induce the DKD phenotype. The hyperglycemic phenotype in DKD patients typically takes years to manifest, thus the use of higher levels or cytokine stimulation in an experimental model may be necessary to observe a phenotype within a reasonable time frame. The levels we tested are aligned with previous studies of *in vitro* diabetic vasculopathy and DKD-sensitized COVID-19 in iPS cell-derived organoids, which utilized 75 mM and 25 mM glucose, respectively.^18,20^

Organoids also exhibit metabolic immaturity compared to kidney tubules *in vivo,* which could contribute to the need for high glucose concentrations to elicit phenotypes^60^. Of note, kidney organoids are typically differentiated in 11 mM glucose, whereas normal glucose levels (3.9 - 5.5 mM) may be too low for healthy maintenance *in vitro*, although more studies are needed to determine this effect.^14^

Another limitation is that organoids derived from iPS cells *in vitro* lack a functional vascular system, and therefore, the effect of glucose on podocyte filtration or tubular reabsorption cannot yet be assessed with this system. Although kidney organoids also contain endothelial cells derived from iPS,^14^ such cells have been previously studied in vascular organoids,^20^ thus we have focused on nephron epithelial cell types in this study. These proof of concept studies were conducted with cells originating from patients who did not have diabetes. Given the wide variability between iPS cell lines from different individuals, a convincing study of DKD in patient-derived cohorts would require well-matched groups of progressors versus controls, which would be interesting to conduct in the context of high glucose.

Finally, further work is needed to fully understand the cascade of molecular events that lead from high glucose exposure to epithelial cell detachment. While our findings suggest that inhibitors of MIF, TNF-alpha, and MEK can be used to intervene in this process, careful consideration and preclinical studies are needed to determine which of these treatment strategies is likely to be most effective, and how they can best be deployed to prevent podocyte spreading and associated kidney dysfunction. The paradigm suggested by these studies, in which high glucose triggers a tissue-intrinsic inflammatory response that leads to epithelial cell detachment, is worthy of further exploration in the kidneys as well as other epithelial organs and organoids, in developing a more detailed understanding of DKD and its therapeutic strategies.

## METHODS

### Stem Cell Culture

WTC11 iPS cells (Coriell GM25256), derived from a Japanese male donor (gift of Dr. Bruce Conklin, Gladstone Institute), and H9 human embryonic stem cells were used to differentiate kidney organoids. WTC11 cells were cultured with 2mL of mTeSR1 (Stem Cell Technologies) supplemented with 1% penicillin/streptomycin (Gibco) in 6-well plates (Falcon) coated with 0.5% Geltrex (Gibco). Cells were incubated at 37 C⁰ and 5% CO2 and passaged using ReLeSR (STEMCELL Technologies).

### Kidney Organoid Differentiation

WTC11 iPS and H9 ES cells were differentiated in kidney organoids following the protocol previously published by our lab. In summary, tissue plates were precoated with 0.5% GelTrex for 11 at 37C⁰. Afterwards, cells were dissociated with Accutase (STEMCELL Technologies) and 100-400 (WTC-11) or 800-1.2k (H9) cells per well were seeded into 96-well (Greiner Bio-One), where else 1-2k cells per well were seeded on or 24-well plates (Falcon), in supplemented with 10 μM Y-27632 ROCK Inhibitor (STEMCELL Technologies). On the following morning single cells were sandwiched with 40% of Matrigel (Corning) in mTeSR1, and media was replaced the following day. On day 3 spheroids were treated with 12 µM CHIR (Stemgent) (WTC-11) or 10 µM CHIR + 10 ng/ml Noggin (PeproTech) in aRPMI (Thermo Fisher Scientific) for 36 to 40 hours. Afterwards media was replaced with RB (aRPMI + 1X Glutamax + 1X B27 Supplement, all from Thermo Fisher Scientific) and replaced every 3 days thereafter until maturation (days 18-21).

### Glucose treatment

Between days 18 and 21, organoids were switched to a novel media formulation with several glucose concentrations (Experimental Day 0). RPMI formulation without glucose (Thermo Fisher Scientific) was mixed with D-Glucose (Sigma-Aldrich) to obtain concentrations of 5.5, 11, 22, 33, 100 mM. Media was afterwards supplemented with 1X Glutamax + 1X B27 Supplement (all from Thermo Fisher Scientific). To test the effect of the cytokines we added 1 ng/ml of TNF-alpha (PeproTech), or 10ng / ml of MIF (PeproTech) or 10ng / ml of TGF-beta (PeproTech) to the media mixed with D-Glucose. Media was then replaced and sampled after three days.

### Inhibitors treatment

Several inhibitors were separately added to the hyperglycemic condition of 33 mM glucose. Inhibitors treatment was performed in parallel with the glucose treatment, six days of treatment started between days 18 and 21 of organoid differentiation. Inhibitors used included ISO-1 (Sigma Aldrich) MIF inhibitor at concentrations: 100, 200, 500 μM; etanercept (Sigma Aldrich) TNF-alpha inhibitor at concentrations: 0.1, 0.5, 1 μg/μl; PD98059 (Stemcell technologies) MEK/ERK inhibitor at working solution of 25 μM; and SB203580 (Selleck chemicals) MAPK inhibitor at working concentration 15 μM. All the inhibitors were delivered with 0.1% DMSO (Corning) vehicle, and stock solution was stored at −20 ℃, except for etanercept which was dissolved in water, and stock solution was stored at −80 ℃.

### Immunochemistry

Kidney organoids were fixed in equal volumes of culture media and 8% PFA (4% working solution) for 15 minutes at room temperature. Afterwards, organoids were washed three times in PBS for 5 minutes and stored at 4 ℃ until use. Samples were blocked for 1h at room temperature with blocking solution (5% donkey serum (Millipore)/0.3% Triton X-100/PBS). Afterwards, primary antibodies were diluted in antibody dilution buffer (3% BSA (Fisher)/ 0.3% Triton X-100/PBS) and incubated overnight at 4 ℃. Primary antibody used: LTL (Vector Laboratories) specie: biotin 1/500 dilution, ECAD (Abcam) specie: rat 1/300 dilution, PODXL (Invitrogen) specie: goat 1/500 dilution, ZO1 (Thermo Fisher Scientific) monoclonal 1/100 dilution. Primary antibodies were washed three times with PBS for 5 minutes; afterwards secondary antibodies (Invitrogen) were diluted 1/1000 in antibody dilution buffer with 1/40 Hoechst 33342 Trihydrochloride trihydrate (Invitrogen) for 1h at room temperature. Samples were then washed and stored in PBS and protected from light to avoid photobleaching. Wild field whole well images 4x images were taken using an Olympus IX83 microscope equipped with a disk spinning unit (DSU, Evident) and cellSense software. Instead 20x confocal images were taken with Nikon Live-Cell Inverted Wide field/Spinning Disk Confocal with a slice / 1μm.

### Time-lapse

kidney organoids were differentiated from GFP-PODXL conjugated WTC11 cells. The same organoid was imaged before glucose treatment, 3 hours after treatment and every 24 hours thereafter for 6 days. Images were taken with an Olympus IX83 microscope, organoid positions were recorded at time zero and used to track the same organoid during the experimental time.

### LDH

Assay was performed using LDH Glo Cytotoxicity assay (Promega) following the user manual protocol. Briefly, the reagent mix was made by mixing the enzyme mix and reductase at 200:1 ratio. Afterwards, 25 μl of reagent mix was added to 25 μl of culture media (collected at each time point and stored −80 ℃) on a 96 well plate (Greiner Bio-One) and incubated for 1h at room temperature. Luminescence was recorded with excitation of 560 nm and emission of 590 nm. Negative control was obtained with black media and positive control by adding 20% DMSO to the organoid alongside glucose treatment.

### Live/dead

Assay was performed by adding growing media to each well with 1/1500 of Calcein AM (live staining, Invitrogen) and 1/1500 Propidium Iodide (dead staining, Invitrogen). The plate was incubated for 20 min at 37 ℃ and immediately after imaged using an Olympus IX83 microscope.

### scRNA-seq

Kidney organoids were differentiated on a 24 well-plate, between days 18 and 21 organoids were treated with a high glucose condition of 33 mM of glucose vs a control condition of 11 mM of glucose. After six days of glucose treatment organoids were dissociated with 1 ml of trypLE (Thermo Scientific) for 15 minutes. Cells were helped to dissociate by gentling pipetting with a p1000 every 5 minutes during the trypLE incubation. Single cells were spun down and resuspended in 1 ml of media with 1% BSA and then filtered using a 10 μm cell strainer.

Cells were resuspended at a concentration of 1000 cellsi/mL in 1X PBS with 1%BSA with 8300 cells being loaded for subsequent 10X Genomics Chromium single cell RNAseq preparation. Single cell droplet libraries were generated in the 10X Genomics Chromium controller according to the manufacturer’s instructions in the Chromium Single Cell 3’ Reagent Kit v.3.1 with Dual Indexing. Additional components used for library preparation include the Chromium Single Cell 3’ Library and Gel Bead Kit v.3.1 (PN-2000164) and the Chromium Single Cell 3’ Chip G kit v.3.1 (PN-120236). Libraries were prepared according to the manufacturer’s instructions using the Chromium Single Cell 3’ Library and Gel Bead Kit v.3.1 (PN-2000164) and Dual Index Plate TT Set A (PN-1000215). Final libraries were pooled at 2nM and sequenced on an Illumina NextSeq2000. Sequencing parameters were selected according to the Chromium Single Cell v.3.1 specifications. Libraries were sequenced to a mean read depth of at least 20,000 total aligned reads per cell.

### scRNA-seq analysis

23,855 cells were sequenced, sequencing base calls were converted to FASTQ files demultiplexed by sample indexes using Illumina’s BaseSpace DRAGEN Analysis Application v1.2.1. FASTQ files were uploaded to 10x Genomics’ Cloud Analysis platform and processed using the Cell Ranger Count Pipeline v7.0.1. Resultant feature/cell matrix (filtered) files were imported to Partek Flow (Illumina) for quality filtering (cells with UMI < 40,000, genes < 8,500, mitochondrial counts < 10%, and ribosomal counts < 40% were kept). Counts were normalized using TMM (Trimmed mean of M-values) for ANOVA differential expression analysis with default parameters. Clusters and sub-clusters were identified using trained Garnett classifiers based on marker genes previously identified for kidney organoid cell types^17,21^. Differentially expressed genes (FDR < 0.05) were subjected to gene set enrichment analysis using the Human_MSigdb_December_01_2022_symbol.gmt version of the MSigdb database (www.gsea-msigdb.org) and the GO bp database (version 2022-12-04). For geneset semantic similarity analysis, enriched GO bp term identifiers were exported and used as inputs for the simplifyEnrichment R package (version 1.14.1). For UMAP representation and cluster analysis data was normalized using SCTransform and principal component analysis (PCA) was performed on the output SC scaled data followed by UMAP (Uniform Manifold Approximation and Projection) dimension reduction or graphed-based clustering under default settings. The new dataset is available on <u>Zenodo</u> through a restricted link.

### Western blotting

After six days of glucose treatment, kidney organoids were lysed with RIPA buffer containing protease and phosphatase inhibitors (Roche). Protein concentration was determined using a Pierce BCA protein assay kit. 50 μg of total protein was separated in a 4–20% acrylamide gel (Bio-Rad) and transferred onto a PDVF membrane using standard procedures. 5% milk was used as a blocking agent antibody dilution buffer. MIF (Rabbit mAb #87501, Cell Signaling) 1:100 and GAPDH Rabbit mAb #14C10, Cell Signaling) 1:2500.

### qPCR

after 6 days of glucose exposure, kidney organoids were washed once with DPBS and incubated with 500 μL per well of Trizol reagent (Invitrogen) for 30 minutes at room temperature. Three wells from a 24 well plate were combined to obtain each sample. RNA was purified using miRNA Isolation Kits (Qiagen). cDNA was synthesized using SuperScript IV Reverse Transcriptase per the manufacturer’s instructions (Thermo Fisher Scientific). Real-time qRT-PCR was performed with PowerUp SYBR Green Master Mix and the following forward and reverse primers: TNF: forward AAGAGATGTGGCAAGAGA, reverse TGTTCATTCATTCATTCATTCAT, β-actin: forward GCGAGAAGATGACCCAGATCAT, reverse GGATCTTCATGAGGTAGTCAGTC. 2^-ddc^ Calculation was used to measure variations in mRNA fold change, beta-actin was used as the housekeeping gene.

### Image analysis

To avoid adding bias during imaging and post-processing, experiments were performed with identical staining and imaging conditions and following homogeneous workflow. Images were then processed using semi-automatic programs written in-house using Fiji (ImageJ), for the image analysis part, and MATLAB or Microsoft Excel to post-process or organize the data. **Organoid remaining**: kidney organoids are counted in Fiji using the *cell counter* plugin from 4x whole-well bright-field images. Organoids were counted at days 0, 3 and 6. Day six immunofluorescence whole well images were used as reference checkpoints to confirm the actual presence of an organoid in the area of interest. We defined as an organoid a structure showing simultaneously signals from the three-nephron marker considered (ECAD, PODXL and LTL). Afterwards days 3 and 6 data were normalized to day 0 the percentage of remaining organoids in each well displayed. **Organoid area and channel intensity**: analysis performed 4x whole-well immunofluorescence images. Kidney organoids were manually circled, being extremely careful to follow the organoid border, afterwards it was calculated the ROI (region of interest) area and the intensity value for each channel (ECAD, PODXL, LTL). Quantification of fluorescent signals is presented as a normalized relative metric after subtraction of the background signal. The background was defined as the average minimum pixel intensity across all images within a given set, afterwards the intensity was normalized to the control condition of 11mM glucose. **Live/Dead:** the percentage of live on dead areas was calculated by dividing the total area from the green channel against the total area from the red channel from whole-well 4x images. **ECAD, PODXL, LTL area calculation**: 20x confocal images were collapsed on a single image using a z projection, then the area of each channel was calculated separately. Afterwards, as shown in figure 2-B, each channel area was divided into *intact and detached area*s, based on a set threshold value, equal across all the experiment, to differentiate the area of the main organoid body (*intact*) identified as part of the organoid comprising the main body, where the three nephron markers are structured and interconnected, vs the area of the single cells/ cluster of cells surrounding the main organoid body and detached from it (*detached*). To evaluate the good functioning of the process we overlaid the area of each channel on the original images and each analysis was afterwards validated. The custom code used for this analysis is provided in the Supplement.

### Statistical

Analysis was performed using GraphPAD Prism, each data point represents either a single well or a single organoid, data are graphs showing average and standard error (SEM). The number of organoids analysed and the experimental number is specified in the figure legends. Data were analyzed using one-way ANOVA and Fisher’s LSD post-hoc testing.

### Bulk RNASeq bioinformatic analysis

For the bulk RNASeq data from Fan et al 2019, we reanalyzed the data at our end downloading the available data from GSE142025. The quality control of this data was done using MultiQC version 1.8. All the RNASeq samples were aligned using GRCh38 reference genome using bulk RNASeq workflow in Seven Bridges using SBG Create Expression Matrix CWL1.0 workflow. Alignment and quantification were done using STAR. Gene symbol mapping was performed using GRCh38ERCC.ensembl99 version. The normalized data were checked for variance contribution and outlier detection using principal component analysis (PCA) function available under DESeq2 package with all available phenotypic covariates in the study design. Visualization for all explanatory phenotypic variables for variance explanation was performed using *ggplot2* function. Subsequent downstream comparative differential expression analysis using DESeq2 to summarize transcript-level estimates for gene-level analysis with covariate of interest in R version 4.2 and BioC version 3.15. For all differential expression comparison statistical thresholds were set as p-adjusted ≤ 0.05 and log2FoldChang +/- 0.8. The comparative assessment of overlapping differentially expression genes across different pairwise comparative groups was done using Venny 2.0 webtool available at (https://bioinfogp.cnb.csic.es/tools/venny/index2.0.2.html).

### Quantification and statistical analysis

Statistical analysis was performed using GraphPad Prism (La Jolla, CA). For each experiment, the number of independent experiments from which the data are pulled and the number of organoids used (*n*) are indicated in the legend. Data are either presented as bar plots with mean ± SEM or an xy plot with SEM indicated at each data point. Data were analyzed using t-tests or one-way ANOVA, and were not corrected for multiple comparisons (Fisher’s LSD test).

### Randomization and Exclusion Criteria

To reduce bias and control for technical variability and batch-to-batch effect, for each experiment, organoids were pulled from different wells for each condition. Each condition was randomly assigned to each well, and the organoids analyzed were randomly selected. Image analysis was performed using in-house code to avoid bias during analysis. Parameters during image acquisition and analysis were kept constant across all experiments, and all conditions were run in parallel. Each organoid was assigned a unique identifier to link each organoid to the specific experiment number, plate, and well. Experiments were excluded if differentiation was unsuccessful (nephron markers not properly defined). Outliers were removed only if the differentiation or staining for the specific organoid was unsuccessful (nephron markers not properly defined or background detected).

### Data Aviability

All primary data supporting the findings of this study are included in the main text and Supplementary Information. Due to the large size and complexity of the raw and processed datasets, public sharing is not feasible. Key data points are visualized as individual dots in the figures throughout the paper and Supplement. Full datasets, including raw data and statistical analyses, are available from the corresponding author upon reasonable request. The scRNAseq dataset is available on Zenodo through a restricted link.

## AUTHOR CONTRIBUTIONS

BSF and AK conceived the project. BSF, AK and GS designed the experiments. GS, MCR, and HHW performed experiments. GS, MCR, VD, HHW, AK, and BSF analyzed the data. GS, HHW, MCR, AK, and BSF wrote the paper.

## ACKNOWLEDGEMENTS

We thank Jonathan Himmelfarb, Stuart Shankland, Sienna Li, and members of the Freedman lab (UW), for funding support, technical support, preliminary experiments, and helpful discussions. Illustrations were created with BioRender.com. Studies were supported by a Novo Nordisk Research Award (BSF), NASA Proposal #21-3DTMPS_2-0001, Department of Defense Award W81XWH2110007, NIH Awards U01DK127553 (BSF), UH3TR003288 (BSF), UC2DK126006 (BSF), and U2CTR004867 (B.S.F.); UW Institute for Stem Cells and Regenerative Medicine Genomics Award (BSF) and Fellowship (GS).

---

**Supplementary Figure S1, related to Figure 1. High glucose induces morphological changes in kidney organoids.** A) Time-lapse phase contrast images showing the progression of the whole plate over six days of culture under four glucose concentrations. B) Quantification of LDH release in culture media between days three and six (mean ± stderr, n ≥ 20 wells per condition pooled from 7 independent experiments)

**Supplementary Figure S3, related to Figure 3. TNF-alpha and TGF-beta sensitize organoids to high glucose.** A) Time-lapse phase contrast images showing the progression of whole wells over six days of culture in four glucose concentrations. B) Quantification of PODXL intact vs detached areas and C-E) PODXL, LTL, ECAD intact areas. The dataset is identical to that analyzed in the main figure, but all data points are shown on the graph including outliers (mean ± stderr, n ≥ 21 organoids per condition pooled from 6 independent experiments; ***P<0.00, **P<0.01, *P<0.05 by one way ANOVA). F) Representative confocal immunofluorescence images showing a single optical section (20x magnification) for each condition. G) Quantification of LDH release in culture media between days three and six (mean ± stderr, n ≥ 19 wells per condition pooled from 6 independent experiments). H) Quantification of PODXL intact vs detached areas and PODXL, ECAD intact areas (mean ± stderr, n ≥ 14 organoids per condition, pooled n ≥ 3 independent experiments. The dataset is identical to that analyzed in the main figure, but all data points are shown on the graph, including outliers. ***P<0.001, **P<0.01, *P<0.05 by one-way ANOVA). I) Quantification of PODXL intact vs detached areas and PODXL and ECAD intact areas (mean ± stderr, n ≥ 14 organoids per condition, pooled from 3 independent experiments. The dataset is identical to that analyzed in the main figure, but all data points are shown on the graph, including outliers ***P<0.001, **P<0.01, *P<0.05 by one-way ANOVA.

**Supplementary Figure S4, related to Figure 4. scRNA-seq analysis reveals high glucose induces cytokine and signaling pathways.** A) Graph showing expression of characteristic genes for each sub-cluster. B) UMAP showing total clusters identified in scRNA-seq analysis. C) Graph indicating 11 vs 33 mM distribution for each cluster of the entire dataset. D) Graph showing expression of characteristic genes for each sub-cluster. E) Graph indicating 11 vs 33 mM distribution for each sub-cluster.

**Supplementary Figure S5, related to Figure 4. scRNA-seq analysis reveals that high glucose induces cytokine and signaling pathways.** A) GO term semantic similarity for the total of podocyte and proximal tubule sub-clusters. B) Graph indicating up- and downregulated pathways with MsigDB Hallmark pathways analysis in the proximal tubule sub-cluster. C) Graph indicating downregulated pathways with MsigDB Hallmark pathways analysis in the podocyte sub-cluster. D) Graph indicating up- and downregulated pathways with KEGG pathways analysis for the full epithelial cluster. E) Graph indicating downregulated pathways with KEGG pathways analysis in the proximal tubule sub-cluster. F) Volcano plot showing up- and down-regulated genes in 33 vs. 11 mM glucose in all clusters combined, with cutoffs of fold change > 1.5 and P value < 0.05. G) Venn diagram showing numbers of genes in each category and overlap from comparison between up and down-regulated genes (FDR<0.05 and log_2_FC ± 0.8) in glucose-treated organoids and a human dataset of kidney biopsy samples from patients with advanced DKD, with H) distribution of each gene in a correlation matrix. I) Comparison of P values for genes in common between the organoids and the patients with advanced DKD.

**Supplementary Figure S6, related to Figures 5 and 6. Pharmacological inhibitors protect organoids from the effects of high glucose.** A) Whole uncropped MIF immunoblot from the main figure. B-D) Quantification of PODXL intact vs detached areas, E) Representative confocal immunofluorescence images showing a single optical section (20x magnification) comparing 11 mM glucose, 33 mM, and 33 mM glucose + Etanercept for WA09 ES cells. F) Quantification of PODXL intact vs detached areas (mean ± stderr, n ≥ 18 organoids per condition, pooled from 2 independent experiments). G) Quantification of PODXL intact vs detached areas and h) PODXL intact/detached areas ± SB23580 (MAPK Inhibitor) and ± PD98059 (MEK/ERK inhibitor). The dataset is identical to that analyzed in the main figure, but all data points are shown on the graph including outliers (mean ± stderr, n ≥ 19 organoids per condition pooled from 3 independent experiments. ***P<0.00, **P<0.01, *P<0.05 by one way ANOVA). I-J) Quantification of PODXL area. Dataset is identical to that analyzed in the main figure, but all datapoints are shown on the graph, including outliers (mean ± stderr, n ≥ 12 organoids per condition pooled from 4 independent experiments; ***P<0.00, **P<0.01, *P<0.05 by one-way ANOVA).

