## Supplementary Figures for "Elevated glucose in kidney organoids induces tissue-intrinsic inflammation driving epithelial detachment"

### SUPPLEMENTARY FIGURE 1

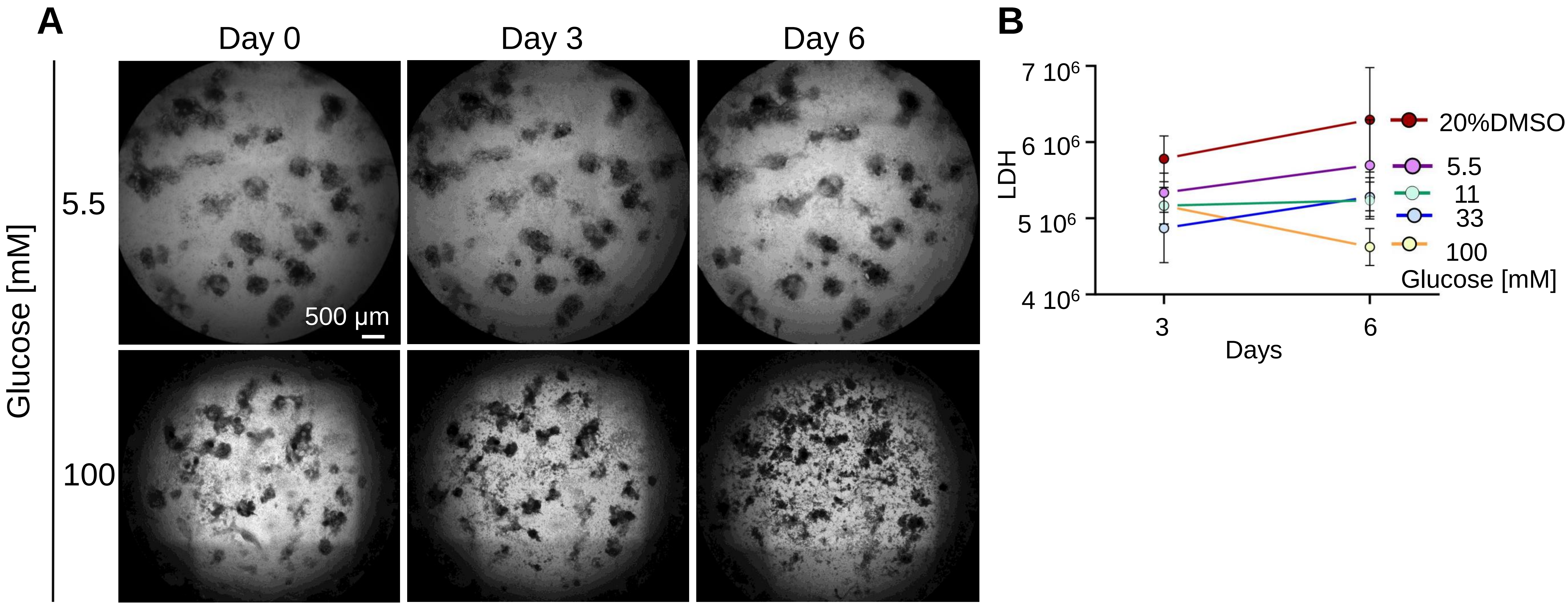


**Supplementary Figure S1, related to Figure 1. High glucose induces morphological changes in kidney organoids. A)** Time-lapse phase contrast images showing the progression of the whole plate over six days of culture under four glucose concentrations. **B**) Quantification of LDH release in culture media between days three and six (mean ± stderr, n ≥ 20 wells per condition pooled from 7 independent experiments)

### SUPPLEMENTARY FIGURE 2

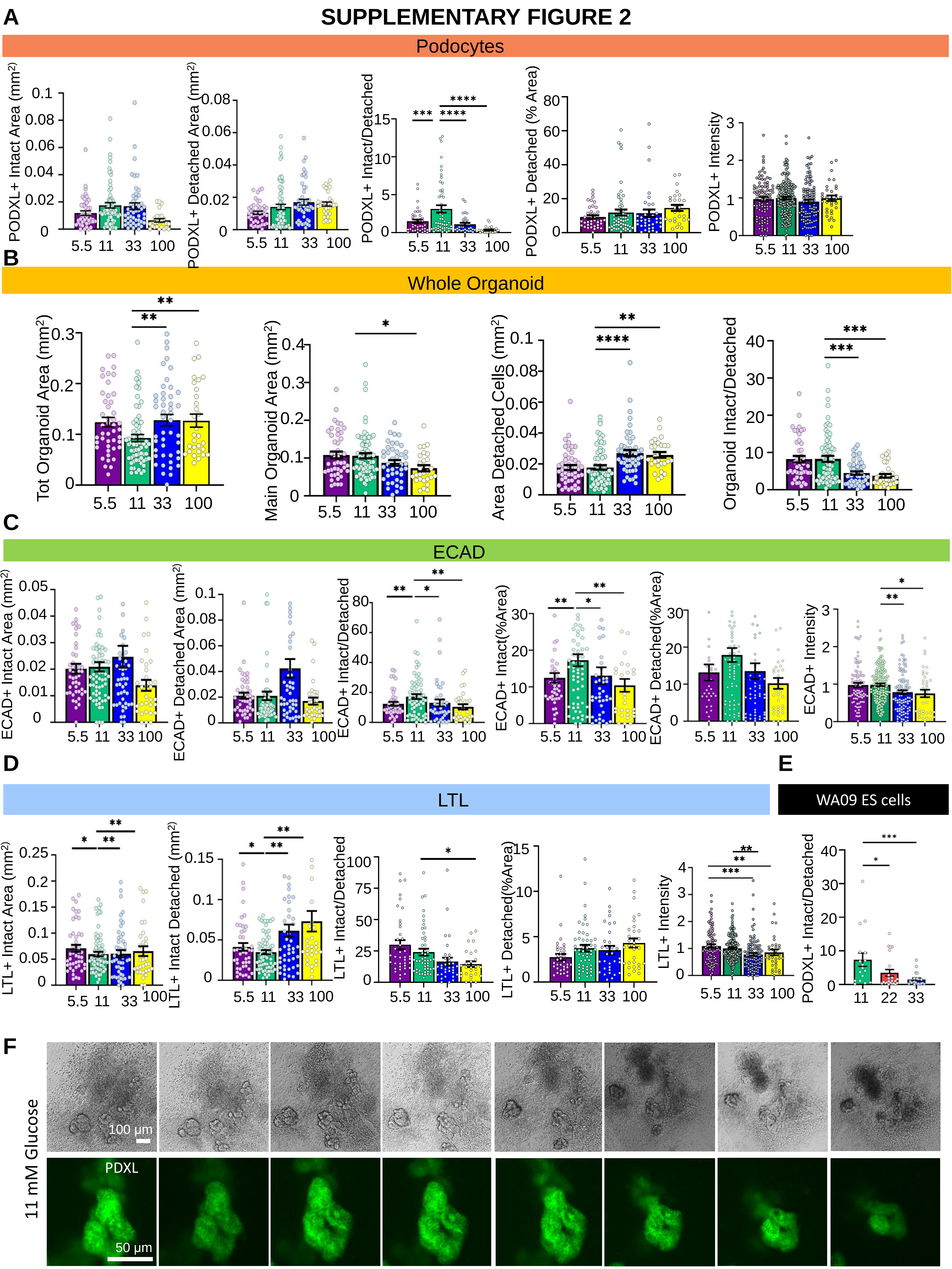

**Supplementary Figure S2, related to Figure 2. High glucose induces the detachment of organoid nephron segments. A)** Quantification of size metrics and ratios for PODXL **P<0.01, *P<0.05 by one-way ANOVA). **B**) Quantification of whole organoid, **C)** ECAD, and **D)** LTL. All data points are graphed for conditions in which the outliers are not graphed in the main figures. For size, statistics, and datasets are the same as in the main figures. Quantification is also shown for intensity of these segments (mean ± stderr, n ≥ 40 organoids per condition pooled from 6 independent experiments; ***P<0.00, **P<0.01, *P<0.05 by one-way ANOVA). **E)** PODXL intact/detached area in WA09 ES cells (mean ± stderr, n ≥ 18 organoids per condition pooled from 2 independent experiments; ***P<0.00, **P<0.01, *P<0.05 by one-way ANOVA). **F)** Representative time-lapse confocal immunofluorescence and phase contrast images showing single optical sections (20x magnification) of PODXL-GFP organoids.

### SUPPLEMENTARY FIGURE 3

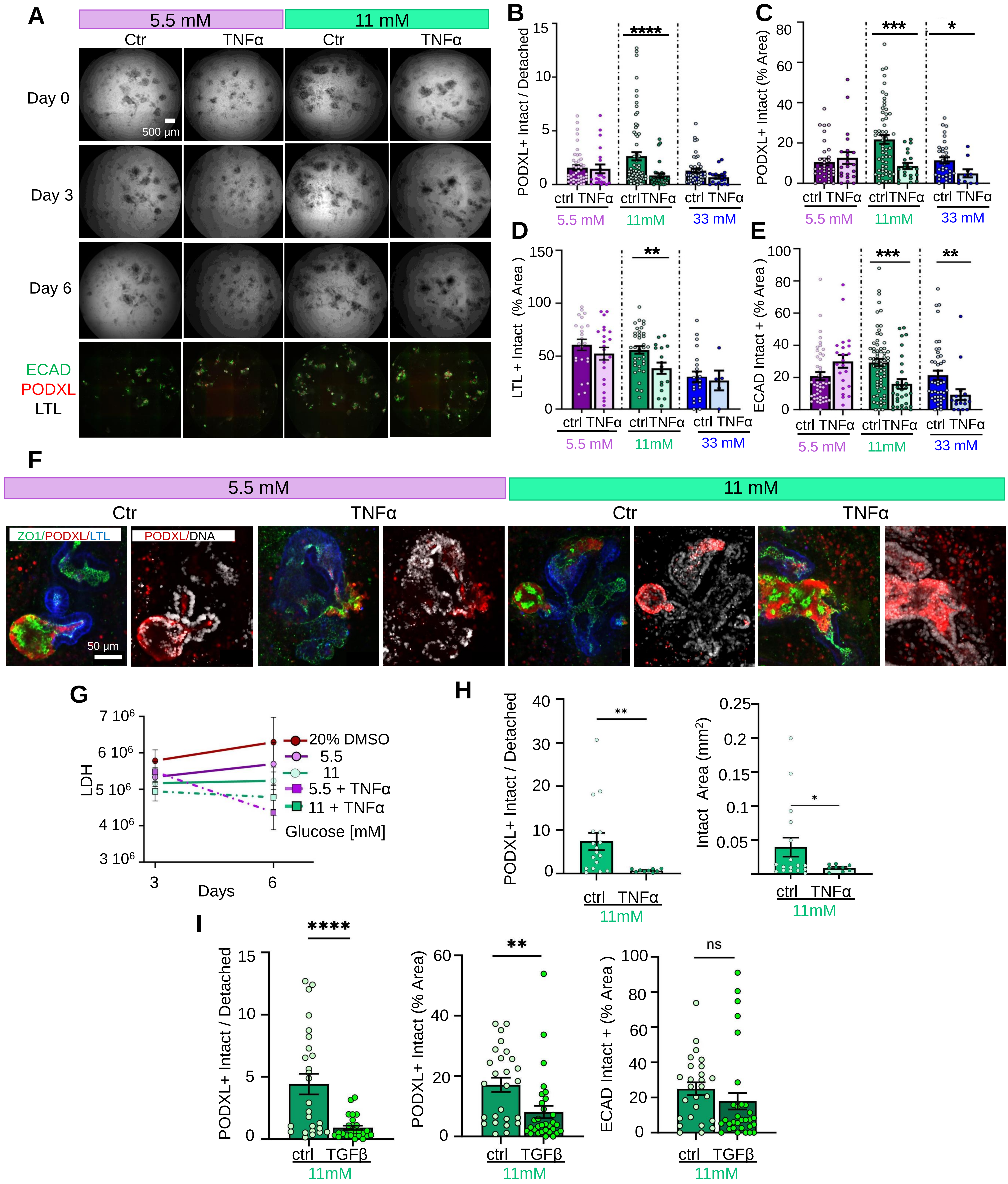

**Supplementary Figure S3, related to Figure 3. TNF-alpha and TGF-beta sensitize organoids to high glucose. A)** Time-lapse phase contrast images showing the progression of whole wells over six days of culture in four glucose concentrations. **B)** Quantification of PODXL intact vs detached areas and **C-E)** PODXL, LTL, ECAD intact areas. The dataset is identical to that analyzed in the main figure, but all data points are shown on the graph including outliers (mean ± stderr, n ≥ 21 organoids per condition pooled from 6 independent experiments; ***P<0.00, **P<0.01, *P<0.05 by one way ANOVA). **F)** Representative confocal immunofluorescence images showing a single optical section (20x magnification) for each condition. **G)** Quantification of LDH release in culture media between days three and six (mean ± stderr, n ≥ 19 wells per condition pooled from 6 independent experiments). **H**) Quantification of PODXL intact vs detached areas and PODXL, ECAD intact areas (mean ± stderr, n ≥ 14 organoids per condition, pooled n ≥ 3 independent experiments. The dataset is identical to that analyzed in the main figure, but all data points are shown on the graph, including outliers. ***P<0.001, **P<0.01, *P<0.05 by one-way ANOVA). **I)** Quantification of PODXL intact vs detached areas and PODXL and ECAD intact areas (mean ± stderr, n ≥ 14 organoids per condition, pooled from 3 independent experiments. The dataset is identical to that analyzed in the main figure, but all data points are shown on the graph, including outliers ***P<0.001, **P<0.01, *P<0.05 by one-way ANOVA.

### SUPPLEMENTARY FIGURE 4

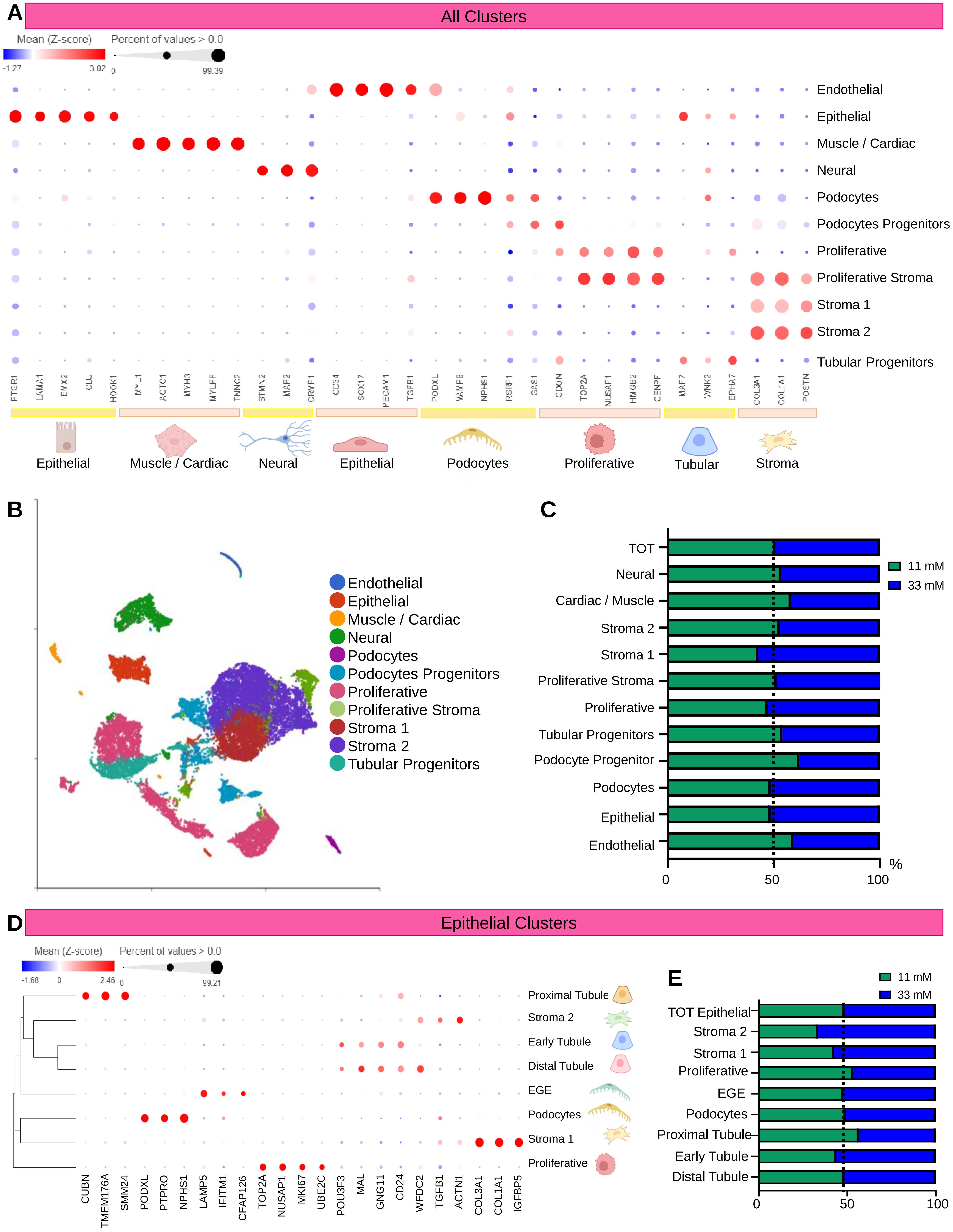

**Supplementary Figure S4, related to Figure 4. scRNA-seq analysis reveals high glucose induces cytokine and signaling pathways**. **A)** Graph showing expression of characteristic genes for each sub-cluster. **B)** UMAP showing total clusters identified in scRNA-seq analysis. **C)** Graph indicating 11 vs 33 mM distribution for each cluster of the entire dataset. **D)** Graph showing expression of characteristic genes for each sub-cluster. **E)** Graph indicating 11 vs 33 mM distribution for each sub-cluster.

### SUPPLEMENTARY FIGURE 5

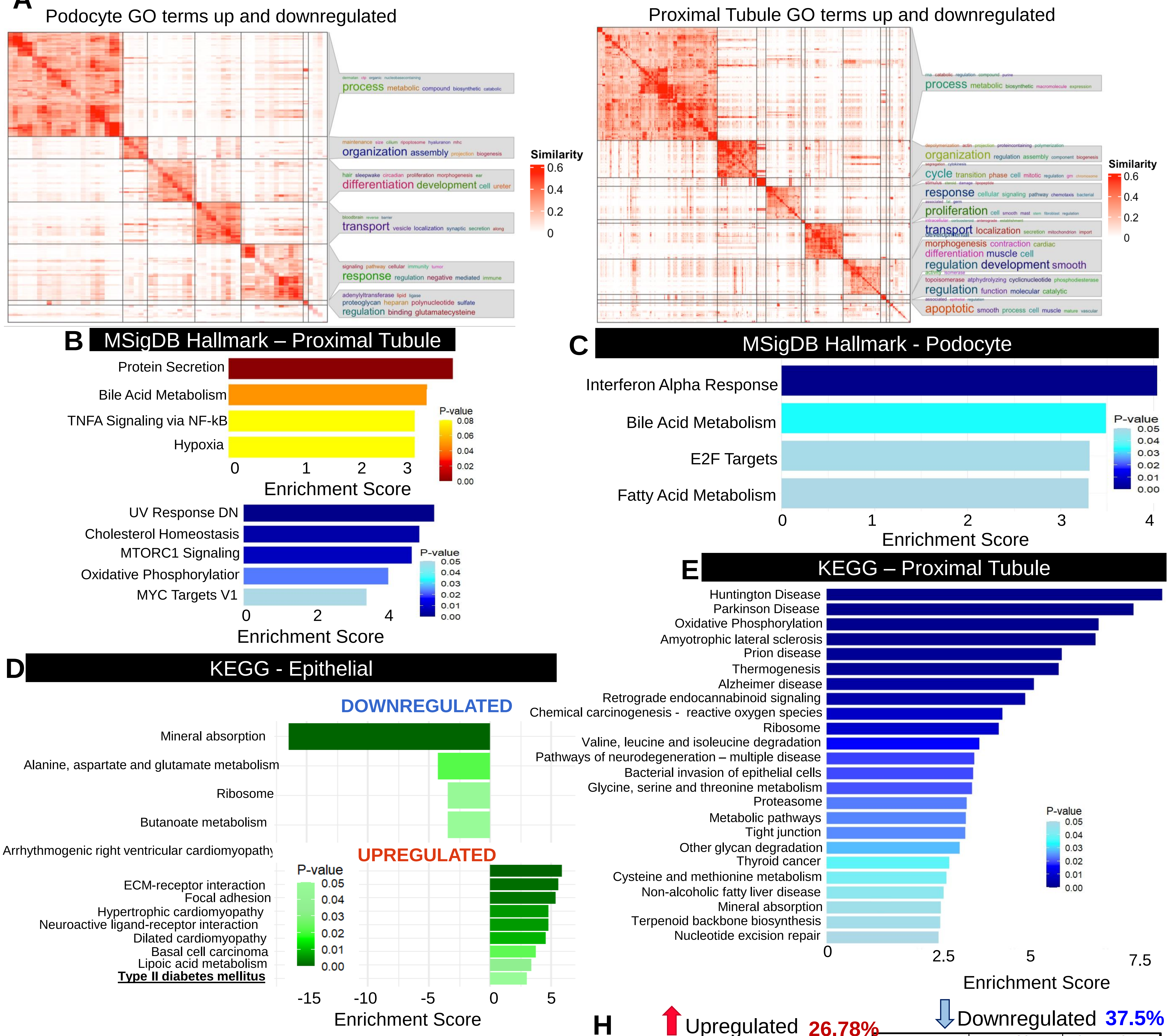


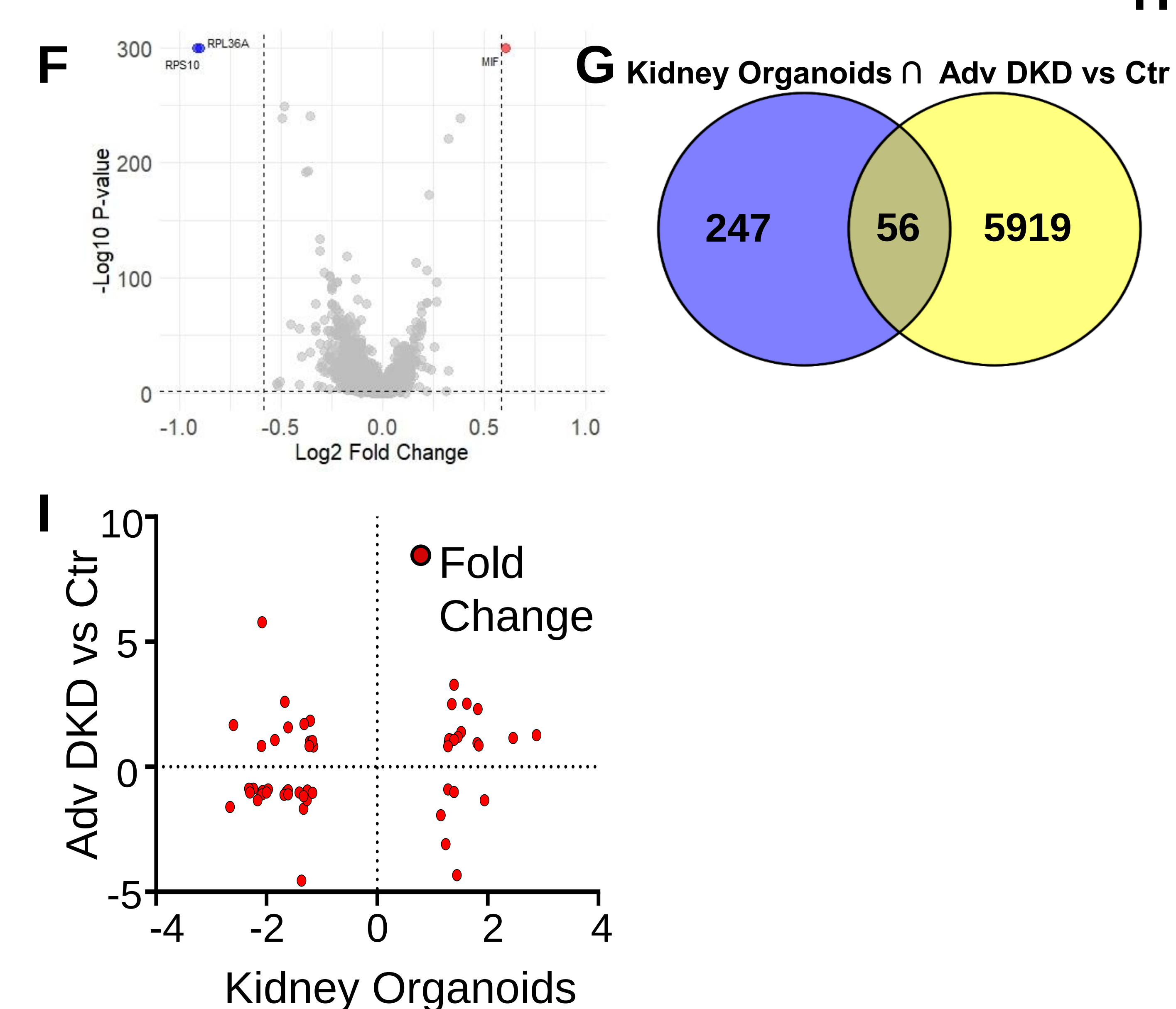


**H**

Upregulated 26.78%

| Gene | Organoid | Advanced DKD |
|---|---|---|
| IFITM1 | 2.88 | 1.2608 |
| KCNIP4 | 1.52 | 1.3857 |
| WFDC2 | 1.35 | 2.5064 |
| SIPA1 | 1.81 | 0.9413 |
| TNFRSF12A | 1.33 | 1.0859 |
| ITGB6 | 1.82 | 2.311 |
| FSTL1 | 1.46 | 1.1886 |
| CRIP1 | 2.46 | 1.1504 |
| FKBP10 | 1.29 | 0.9701 |
| PNOC | 1.39 | 3.2804 |
| ADRA2C | 1.84 | 0.8513 |
| MTHFD1L | 1.3 | 1.1025 |
| TMEM132A | 1.39 | 1.0821 |
| PDLIM4 | 1.28 | 0.8215 |
| LINC01993 | 1.62 | 2.5232 |

Downregulated 37.5%

| Gene | Organoid | Advanced DKD |
|---|---|---|
| ACSM2B | -2.11 | -1.0508 |
| MT-ATP8 | -1.37 | -4.5543 |
| SEC62 | -1.26 | -0.9471 |
| ACSM2A | -2.07 | -0.9609 |
| HMGN5 | -1.33 | -1.6781 |
| GLYAT | -2.08 | -1.1 |
| C11orf54 | -1.41 | -1.031 |
| AGXT2 | -2.24 | -0.8701 |
| FMO1 | -1.64 | -0.9952 |
| INSR | -1.27 | -1.3257 |
| AFP | -2.16 | -1.3414 |
| MT2A | -2.32 | -0.8736 |
| HAO2 | -1.97 | -0.9048 |
| SLC34A1 | -2.30 | -1.0288 |
| SPAG5 | -2.00 | -1.0319 |
| GPHN | -1.17 | -1.0344 |
| AMN | -1.33 | -1.1762 |
| VIL1 | -1.61 | -0.9333 |
| APOC3 | -2.66 | -1.6065 |
| GATM | -1.68 | -1.1168 |
| GLYATL1 | -1.61 | -1.1 |

**Supplementary Figure S5, related to Figure 4. scRNA-seq analysis reveals that high glucose induces cytokine and signaling pathways**. **A)** GO term semantic similarity for the total of podocyte and proximal tubule sub-clusters. **B)** Graph indicating up- and downregulated pathways with MsigDB Hallmark pathways analysis in the proximal tubule sub-cluster. **C)** Graph indicating downregulated pathways with MsigDB Hallmark pathways analysis in the podocyte sub-cluster. **D)** Graph indicating up- and downregulated pathways with KEGG pathways analysis for the full epithelial cluster. **E)** Graph indicating downregulated pathways with KEGG pathways analysis in the proximal tubule sub-cluster. **F)** Volcano plot showing up- and down-regulated genes in 33 vs. 11 mM glucose in all clusters combined, with cutoffs of fold change > 1.5 and P value < 0.05. **G)** Venn diagram showing numbers of genes in each category and overlap from comparison between up and down-regulated genes (FDR<0.05 and $log_2FC \pm 0.8$) in glucose-treated organoids and a human dataset of kidney biopsy samples from patients with advanced DKD, with **H)** distribution of each gene in a correlation matrix. **I)** Comparison of P values for genes in common between the organoids and the patients with advanced DKD.

### SUPPLEMENTARY FIGURE 6

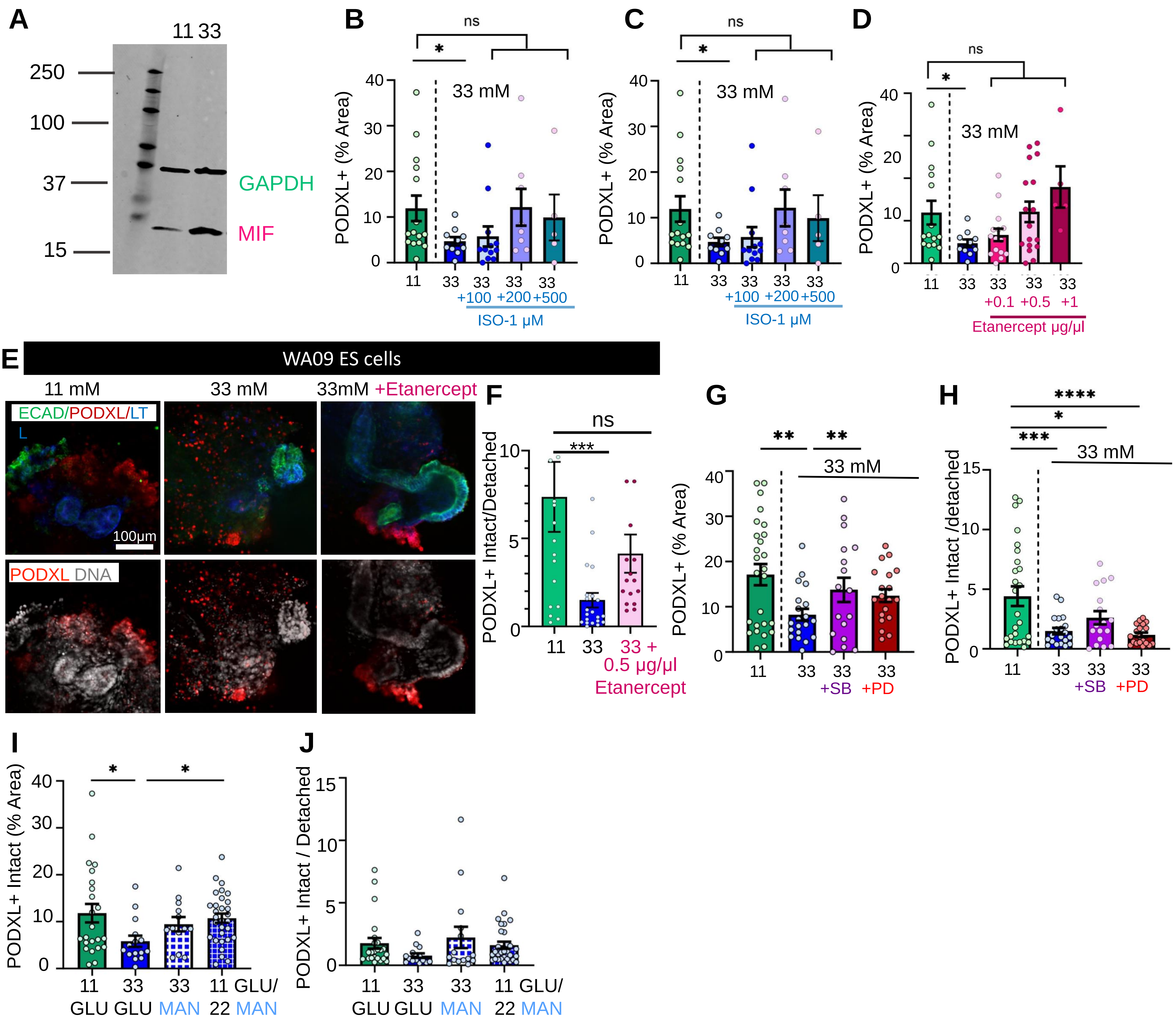


**Supplementary Figure S6, related to Figures 5 and 6. Pharmacological inhibitors protect organoids from the effects of high glucose. A)** Whole uncropped MIF immunoblot from the main figure. **B-D)** Quantification of PODXL intact vs detached areas, **E)** Representative confocal immunofluorescence images showing a single optical section (20x magnification) comparing 11 mM glucose, 33 mM, and 33 mM glucose + Etanercept for WA09 ES cells. **F)** Quantification of PODXL intact vs detached areas (mean ± stderr, n ≥ 18 organoids per condition, pooled from 2 independent experiments). **G)** Quantification of PODXL intact vs detached areas and **H)** PODXL intact/detached areas ± SB23580 (MAPK Inhibitor) and ± PD98059 (MEK/ERK inhibitor). The dataset is identical to that analyzed in the main figure, but all data points are shown on the graph including outliers (mean ± stderr, n ≥ 19 organoids per condition pooled from 3 independent experiments. ***P<0.00, **P<0.01, *P<0.05 by one way ANOVA). **I-J)** Quantification of PODXL area. Dataset is identical to that analyzed in the main figure, but all datapoints are shown on the graph, including outliers (mean ± stderr, n ≥ 12 organoids per condition pooled from 4 independent experiments; ***P<0.00, **P<0.01, *P<0.05 by one-way ANOVA).
